# Prenatal immune priming to *Plasmodium falciparum* is characterized by multi-lineage T-cell responses that are influenced by maternal microchimerism and predict malaria outcomes

**DOI:** 10.64898/2026.10.13.759013

**Authors:** Yonghou Jiang, Florian Bach, John Houck, Marc Carlson, Almahamoudou Mahamar, Santara Gaoussou, Oumar Attaher, Robert Morrison, Sudhir Kumar, Blair Armistead, Irfan Zaidi, Stefan Kappe, Alassane Dicko, Patrick Duffy, Michal Fried, Marion Pepper, Whitney E. Harrington

## Abstract

In malaria endemic settings, the fetus may be exposed to *Plasmodium falciparum* (Pf) antigens leading to prenatal immune primming (PIP). The fetus also acquires a graft of maternal cells in utero, a phenomenon known as maternal microchimerism (MMc). We have previously identified that MMc at birth predicts altered malaria susceptibility in early childhood, and we hypothesized that this association may be mediated via two potential mechanisms. First, the fetus may acquire Pf-specific MMc T-cells that directly respond to postnatal parasitemia. Alternatively, MMc may indirectly influence the quantity or quality of PIP. To investigate this hypothesis, infant cord bloods from 97 Malian primigravidae were screened for MMc using qPCR. A subset of ten cord blood mononuclear cells with variable MMc levels were stimulated with purified Pf merozoite extract and activated T-cells were sorted by flow cytometry for scRNAseq analysis. Transcriptional profile was used to differentiate Pf-responsive versus bystander activated T-cells, and MMc cells were identified using the Cellector algorithm. Among Pf-responsive cells, we identified four key populations: activated CD4 T-cells, ZBTB16+ CD4 T-cells, activated Vδ1+ γδ T-cells, and cytotoxic Vγ9Vδ2+ γδ T-cells. The minority of sorted cells were of MMc origin; however, there was a strong association between the presence of MMc and the quantity and quality of PIP. MMc+ infants had significantly fewer Pf-responsive T-cells, as well as a shifted distribution of these cells with fewer ZBTB16+ CD4 and cytotoxic Vγ9Vδ2+ γδ T-cells. Clinically, MMc was protective from incident parasitemia and febrile malaria, whereas the frequencies of ZBTB16+ CD4 and activated CD4 T-cells were associated with increased risk. Together, these data demonstrate that MMc influences malaria outcomes through regulation of offspring inflammatory responses to Pf.

## INTRODUCTION

*Plasmodium falciparum* (*Pf*) malaria continues be to a leading cause of global morbidity and mortality, with an estimated 249 million cases and 608,000 deaths in 2022 (1). Pregnant women and children are particularly susceptible, and children under five account for approximately 76% of deaths (1). During pregnancy, infected erythrocytes bind to the synciotrophoblast, a condition known as placental malaria, leading to early pregnancy loss, stillbirth, and low birth weight (2, 3). In addition, epidemiologic studies suggest a link between placental malaria in the mother and increased susceptibility to malaria in the infant (4, 5). This association has been hypothesized to be the result of in utero exposure to Pf antigens leading to prenatal immune priming (PIP) and an altered postnatal immune response to malaria.

Supporting the notion of PIP, early studies from malaria endemic settings demonstrated cord blood lymphoproliferation in response to malaria antigens (6, 7). More recent work demonstrated the presence of CD4 T-cells producing Th1 and Th2 cytokines, as well as IL10, following stimulation with erythrocyte surface proteins or whole parasite extract (8–13). The nature and degree of malaria exposure in the pregnancy may influence the quality of the T-cell response in cord blood. For example, in one study cord blood from primigravidae and women with peripheral or placental malaria infection was more likely to contain Pf-responsive T-cells, with nearly half of total cord bloods responding to at least one Pf antigen (12). In a second study, placental malaria at delivery was associated with Pf-specific IL10 secreting T-cells in cord blood, whereas treatment of malaria infection prior to delivery was associated with IFNγ production by T-cells (14). Limited work has demonstrated an association between the presence and quality of cord blood Pf-specific T-cell responses and postnatal antibody responses to malaria blood stage antigens (15), as well as susceptibility to malaria during infancy (16, 17).

While the Pf-specific T-cell responses in cord blood have generally been attributed to fetal soluble antigen exposure leading to PIP (18), early work proposed that these responses may reflect instead “the transfer of maternal lymphocytes… to the foetus” (6). Indeed, we and others have demonstrated that the fetus acquires a graft of maternal cells, a phenomenon known as maternal microchimerism (MMc), which is enriched for T-cells (19, 20). We have further found that both maternal peripheral (21) and placental malaria (22) are associated with increased cord blood MMc. Finally, we have found in a cohort of Tanzanian infants that the presence of cord blood MMc was associated with more malaria infections during infancy but protection from symptoms when infected (22). Murine studies of MMc indicate that in utero acquired maternal T-cells may impact offspring immunity in both antigen-specific (23) and antigen-independent manners (24). Based on these observations, we hypothesized that the fetus acquires maternal Pf-specific T-cells in utero, which may in turn impact offspring susceptibility to malaria via at least two potential mechanisms. First, MMc Pf-specific T-cells may directly respond to parasites acquired postnatally, controlling disease. Alternatively, MMc may indirectly influence the quantity or quality of PIP, “educating” the offspring response.

To investigate this hypothesis, we have leveraged single cell RNA sequencing (scRNAseq) to determine the origin and phenotype of Pf*-*responsive T cells at single cell resolution in Malian cord blood. We additionally determined whether MMc cells may may serve as cellular messengers, influencing the quality of the fetal response, and ultimately susceptibility to malaria during infancy.

## MATERIALS & METHODS

### Cohort & Human Subjects

The cohort for the present study was derived from the Immuno-Epidemiology project conducted in Ouelessebougou, Mali between September 2011 and May 2015 (25). Briefly, 97 maternal-infant pairs were selected based on maternal primi- or secundigravidity (26), rationalizing that there would be the greatest probability of identifying malaria-specific T-cells in the cord blood of these women. Mothers were enrolled and followed throughout the pregnancy with diagnosis of clinical malaria based on symptoms and positive blood smear. Placental malaria was diagnosed at the time of delivery with blood smear of placental blood. Infants were followed across the first year of life with scheduled and walk in visits when clinically unwell; blood smears were taken at both routine and walk-in visits. Incident parasitemia in the infant was defined as a positive blood smear following a negative blood smear or at least 28 days following a prior incident parasitemia. Febrile malaria was defined as reported fever recorded by the clinician in the presence of parasitemia. The protocol and study procedures of the parent cohort were approved by the Institutional Review Board (IRB) of the National Institute of Allergy and Infectious Diseases at the National Institutes of Health and the Ethics Committee of the Faculty of Medicine, Pharmacy and Dentistry at the University of Bamako, Mali. All women provided written informed consent for themselves and their infants. Further immunology studies were approved by the Seattle Children’s IRB (STUDY00002140). In addition, the use of samples from healthy adult donors with and without a prior history of Pf infection from the SCRI CGIDR Biorepository was approved by the Seattle Children’s IRB (STUDY00002048); all individuals provided written informed consent.

### Microchimerism screening

To limit the possibility of maternal blood contamination, cord blood was collected at the time of delivery via cross-clamping of the umbilical cord and direct cannulation of the umbilical vessels. Whole cord blood collected into EDTA tubes was (1) frozen and stored at –70°C, and (2) processed for cord blood mononuclear cells (CBMC) and stored in liquid nitrogen. Genomic DNA was extracted from whole cord and maternal blood using QiaAMP DNA Micro or Mini Kits. Human leukocyte antigen (HLA) and non-HLA typing was conducted for all mothers and infants to identify a maternal-specific allele for each pair, as previously described (22, 27). The maternal-specific allele for each pair was amplified from cord blood using a panel of quantitative PCR assays (qPCR), and MMc level was calculated as previously described using number of genomic equivalents (GE) assessed as the denominator (19).

### Purification of *Plasmodium falciparum* merozoites extract (PfME)

Pf parasites were cultured in RPMI1640 medium supplemented with Type O+ human serum, O+ red blood cells (RBC), and 5% CO_2_ / 5%O_2_ / 90% N_2_. The parasites were synchronized with 5% sorbitol at ring stages and were harvested when parasitemia was more than 12%. Merozoites were purified by filtration of highly synchronous populations of late segmenter stage schizonts through a 1.2µM Acrodic 32-mm syringe filter (Pall) and magnet separation column (MACS, Miltenyi Biotec). Pf merozoite extract (PfME) was obtained by treating the purified merozoite pellet through three cycles of freeze/thaw. Purified uninfected RBC extract (puRBCE) was processed via the same protocol and used as a negative control for assay validation.

### Stimulation of cells with PfME

To identify Pf-specific T cells, we first optimized a T-cell stimulation assay utilizing PfME. PfME was selected as the stimulus as our goal was to capture a broad response against parasite blood stage antigens, while limiting RBC antigen exposure. We optimized our assay to reduce background responses to both media only and puRBCE negative controls, thus increasing the probability that cells expressing activation induced markers (AIM) at either the protein or transcript level reflected responses to Pf antigens rather than to RBC antigens.

At the time of stimulation, cryopreserved control peripheral blood mononuclear cells (PBMC) and CBMC were thawed in a 37°C water bath and washed in warm RPMI1640 containing 20% FBS. After washing and centrifugation, the cell pellet was resuspended in 2mL culture media (RPMI1640 containing 5% human sera) and rested at 37°C overnight. The next morning, 100µL of suspended cells (1-3x10^6^) was seeded into individual wells of a 96 well plate in culture media. 1µL of antihuman CD40 antibody (Miltenyi,100µg/ml) was added into each sample well for 20 minutes at 37°C at a final concentration of 0.5µg/mL.100µL of puRBCE or PfME were then added to negative control and stimulation wells respectively and incubated at 37°C for 18hrs. puRBCE and PfME conditions were run for our adult positive control donor during assay validation (**Fig S1**). To maximize the number of cells sorted for scRNAseq for Malian CBMC, only the PfME stimulation condition was conducted.

### Single cell RNA sequencing

Following stimulation, the cell pellet was washed and incubated with the human FcR blocking solution (Human TruStain FcX, BioLegend) for 10 min at room temperature, followed by staining with an antibody cocktail of fluorescently-labeled antibodies (CD3, CD69) and oligo-conjugated antibodies (CD4, CD8, CD40L, CD137, CCR7 and CD45RO, all BioLegend) (hereafter referred to as “feature barcodes”) and viability dye at 4^0^C in the dark for 30 minutes (**Table 1**). After staining, cells were washed and resuspended in PBS with 2% FBS. Cells were sorted on a BD FACS Melody with collection of the CD3+ CD69+ fraction for further analysis. The sorted cells were stored on ice and concentrated into a small volume (<37.5 µL) before proceeding with 10X Chromium scRNAseq.

**Table 1.**
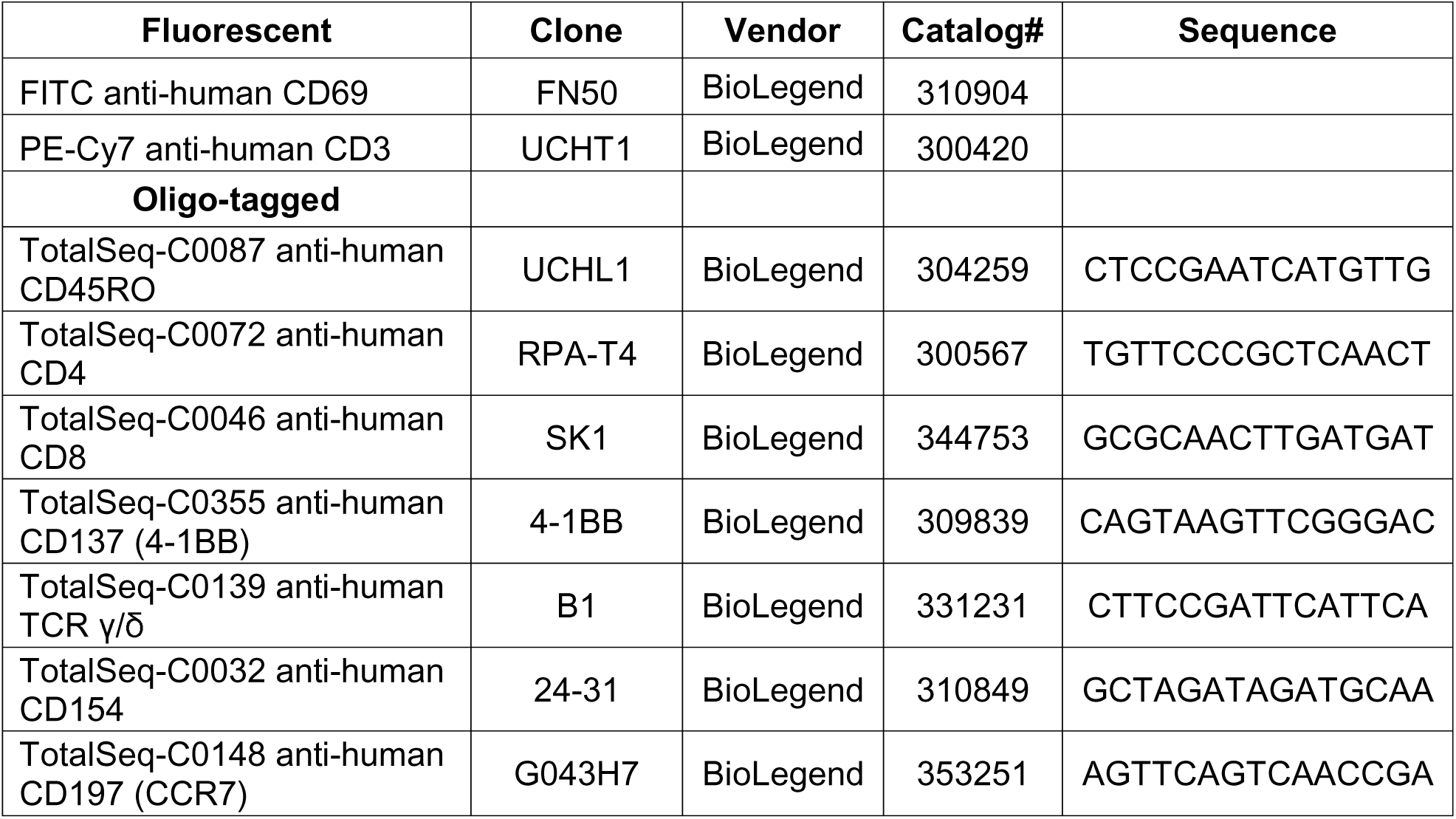
Antibodies used for FACS sorting and scRNAseq.

The range of cell input per sample into the scRNAseq assay was 3,300-15,000. The Chromium Next GEM Single Cell V(D)J Reagent Kit v1.1 and the Chromium Next GEM Single Cell 5’ Reagent Kit v2 (10X Genomics, Pleasanton, CA USA) on the Chromium Controller (10X Genomics, Pleasanton, CA USA) were used following the manufacturer’s recommended protocols. Gene expression (GEX), V(D)J, and feature barcode libraries were generated for each sample. All libraries were assessed to ensure appropriate product size and yield on the Agilent 2100 Bioanalyzer using the Agilent High Sensitivity DNA Assay (Agilent Technologies, Santa Clara, CA USA) following manufacturer’s recommendations and 10X Genomics protocol guidelines. Library yield was further quantified using the NEBNext Library Quant Kit for Illumina® (New England Biolabs, Ipswich, MA USA) following the manufacturer’s recommendations. Libraries were pooled following 10X Genomics guidelines and sequenced with the Illumina NextGen sequencing platform (NovaSeq or NextSeq2000).

### Identification of maternal cells

To identify rare maternal cells in the background of sorted fetal cells, we first utilized the R package Cellector (28) to identify a major and minor population of cells (**Fig S2A**). Cellector is a version of the previously released soupercell (29), refined to better identify minor cell populations from haploidentical individuals, for example in the case of MMc. We then confirmed that cells identified in the minor population in both male and female offspring lacked expression of Y chromosome genes (RPS4Y1, DDX3Y, KDM5D, EIF1AY, ZFY, UTY, and USP9Y) (30) (**Fig S2B**). In addition, in male offspring, we identified any cells that expressed XIST, a long non-coding RNA molecule responsible for Barr body formation (31), as female in origin and thus representing MMc (**Fig S2C**).

### Single cell data quality control and doublet removal

For pre-processing, QC, and base quantification, we used 10X cellranger tools (v7.2.0). The tools ‘cellranger multi’ and ‘cellranger count’, followed by ‘cellranger aggr’ were used to quantify and then to aggregate multiple samples together. Single-cell RNA-seq and antibody capture data from each of the samples was loaded from Cell Ranger raw feature-barcode matrices and processed with Seurat. Each sample was filtered to genes detected in ≥3 cells and cells with ≥200 detected genes. RNA counts were log-normalized (scale factor 10,000) and ADT counts were CLR-normalized. Samples were merged with sample-specific barcode prefixes. Doublets were identified separately for each sample using DoubletFinder. Briefly, each sample was normalized, scaled, and reduced by principal components analysis (PCA), and the number of PCs was selected automatically as the smaller of (i) the first PC at which cumulative variance exceeded 90% and individual variance fell below 5%, and (ii) the last PC preceding a drop in explained variance of <0.1% between successive PCs. Cells were clustered at a resolution of 0.1 to model the proportion of homotypic doublets. The expected multiplet rate was set from the 10x Genomics multiplet-rate table according to the number of recovered cells, and the expected number of doublets was adjusted for the estimated homotypic proportion. Cells classified as doublets were removed.

Remaining cells were filtered to exclude those in the lowest 1% of UMI counts, those outside the 1st-99th percentile range of detected genes, and those in the top 1% of mitochondrial read fraction. Variable features (n = 3,333, vst method) were identified, data were scaled, and PCA was performed. Batch effects between samples were corrected with Harmony (theta = 1, grouping by sample). Cells were clustered on the Harmony embedding, and clusters corresponding to T cells were retained. After clustering, four transcriptionally distinct populations with poor quality metrics (low reads and genes per cell, high mitochondrial percentage) were manually removed. The final remaining T-cell subset was then re-processed with variable feature selection, scaling, PCA, and Harmony integration (theta = 1). The final embedding used the first 30 Harmony dimensions for UMAP and for shared nearest-neighbor graph construction, followed by Louvain clustering at a resolution of 1.0, yielding 15 clusters. After manual merging three populations with largely overlapping phenotypes into one cluster (early activated CD4), 13 populations were taken forward for analysis.

### Identification of Pf-specific T cells

To define activated T-cells, we employed a gene module approach, calculating an activation marker induced (AIM) score based on the expression of the following genes: *CD40LG*, *TNFRSF4*, *TNFRSF9*, *CD69*, *MIR155HG*, *DUSP2*, *IRF4*, *AHR*, *ICOS*, *TNFRSF18*, *NR4A1*, *NR4A2*, and *NR4A3*. The approach was validated using our adult Pf-exposed donor, comparing puRBCE to PfME stimulation, where we identified a clear difference in the frequency of highly activated T-cells, particularly those expressing genes consistent with recent TCR engagement, including Nur77 family members, *CD40LG,* and *ICOS* (**Fig S1D**). To distinguish γδ from ab T cells, we used a similar approach (32), in which a γδ score was calculated using gene expression of all detected gamma and delta V and C genes.

Differential gene expression between donors with and without detectable MMc was assessed by pseudobulk analysis. Donors were classified as MMc-negative (n=4) or MMc-positive (n=6) based on whether any microchimeric cells were detected in the single-cell data prior to quality control. Raw RNA counts were aggregated by donor within each group, and donor-group pseudobulk profiles with fewer than 10 cells were excluded. Differential expression was tested with the Libra R package (run_de) using edgeR with a likelihood ratio test, treating donor as the unit of replication. P values were adjusted for multiple testing using the Benjamini-Hochberg procedure, and genes with an adjusted P value < 0.05 were considered significant. We performed this analysis three ways: comparing gene expression profiles among all T-cells, all cells from the four activated clusters (activated CD4, ZBTB16+ CD4, activated Vδ1, and cytotoxic Vγ9Vδ2) and, lastly, all highly activated T-cells, regardless of cluster. Highly activated T-cells were defined as cells with an AIM signature score ≥ 0.04. 42.5-86.4% of cells in the activated clusters were above this threshold, while 68.7-95.3% of non-activated clusters were below it.

### Statistical analysis

Association between maternal covariates and screened MMc was assessed using logistic regression with adjustment for GE assessed and robust standard errors. Association between screened MMc and risk of parasitemia across the first 18 months of life was assessed within infants followed for at least 4 weeks using a generalized estimating equation (GEE) model with a binomial outcome structure (incident positive or negative blood smear at each study visit), an independent correlation matrix, robust standard errors, and adjustment for placental malaria, number of maternal peripheral malaria infections, age of the child, and GE assessed. Association between screened MMc and risk of febrile malaria was assessed using a similar approach, except that the binomial outcome was defined as presence or absence of fever restricted to parasitemia positive study visits. Association between experimental MMc and frequency of total and subset-specific Pf-responsive T-cells was assessed using t-Tests and multivariate linear regression with adjustment for placental malaria and number of maternal malaria infections in pregnancy. Association between experimental MMc and malaria and non-malarial fever outcomes was assessed using GEE models including all study visits across early childhood and given the low number of events, adjustment was restricted to age of the child for parasitemia and non-malarial fever and no covariates were included for febrile malaria. Finally, association between T-cell subsets and malaria outcomes used the same approach with additional adjustment for the parent frequency of CD69+ cells of CD3+ T cells sorted into 10X scRNAseq.

## RESULTS

### Samples for characterization of the interaction between maternal microchimerism and prenatal immune priming

Mothers included in the present study (n=97) had a median age of 18 (range 15-24). Ninety (93%) were primigravidae and 7 (7%) were secundigravidae; all were primiparous. Eighty-three women (86%) had any peripheral Pf during the pregnancy, with a median of 2 infections (range 0-4). Fifty-three (55%) women had placental malaria at delivery. Median gestational age at delivery was 39 weeks (range 34 to 42 weeks), and 54 (56%) of the infants were female. An informative marker for MMc could be identified in 70 (72%) of the pairs. Of these, 23 (33%) were positive for MMc by qPCR in whole cord blood with levels that ranged from 3 to 2,082 per 10^6^ infant GE. Neither maternal placental nor peripheral malaria was associated with detection of MMc. Among 63 children followed for at least 4 weeks, there were 890 study visits with 56 (6%) incident parasitemia events in the first 18 months of life, among which 33 (59%) were associated with fever. Screening positive for MMc by qPCR was associated with increased risk of incident parasitemia (AOR=2.01, p=0.05) and a trend toward decreased risk of febrile malaria when infected (AOR=0.33, p=0.1), consistent with our prior results from Tanzania (22) (**Fig S3)**.

### Stimulation with PfME induces a polyclonal, multi-lineage T cell response in CBMC

Given the consistent finding that MMc appeared to be associated with early childhood malaria risk, we next sought to investigate the potential mechanisms. We selected ten matched CBMC samples with high cell viability following thawing and overnight rest to move forward to study PIP in detail, including two that screened MMc negative and eight with high MMc levels (greater than 40/10^6^). As described, CBMC were stimulated with PfME and CD69+ CD3+ T cells were sorted for scRNAseq. The frequency of CD69+ cells of parent CD3+ T cells varied from 1.7% to 14.3% across the ten samples. There was no association between maternal placental or peripheral infection and the frequency of CD69+ CD3+ T cells in each sample, however, there was a trend toward fewer sorted cells in screened MMc+ vs MMc-samples (10.3% vs. 4.6%, p=0.1).

After quality control and doublet exclusion, 29,343 cells were retained, with a median of 1,577 genes and 4,226 reads per cell. The median number of cells per CBMC sample was 1,647 (range: 757 to 7,123). Louvain clustering on principal components resolved 15 clusters, which we merged and annotated to 13 final clusters (**Fig 1A**). To label cell subsets, we used canonical lineage and activation markers, metabolic state, as well as TCR chain usage (**Fig 1B, Fig S4**). Because cells were sorted on CD69, the dataset was, as expected, devoid of a resting naive compartment. Instead, it comprised a spectrum of activated conventional CD4⁺ T cells together with regulatory CD4+, CD8⁺, and γδ T cell populations, indicating that PfME stimulation mobilizes a polyclonal, multi-lineage response in cord blood T-cells.

**Figure 1.**
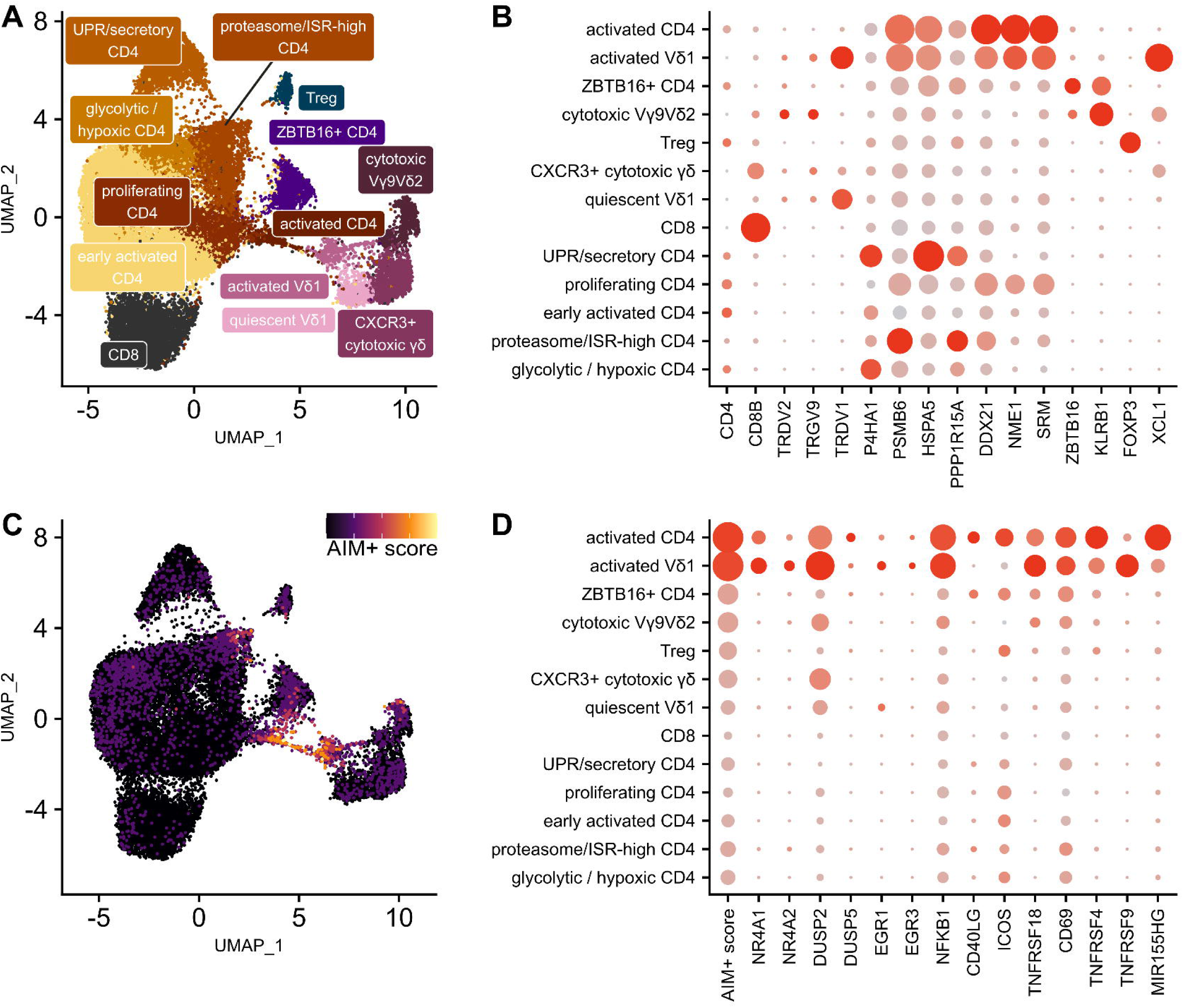
scRNAseq of CD69+ T cells stimulated with PfME reveals diverse CD4 and γδ phenotypes. A). UMAP of all T cells in the dataset, colored by cluster. B) Abbreviated DotPlot showing key markers of lineage and cell state that distinguish each cluster. C) UMAP showing T cells, colored by intensity of our activation marker-derived AIM+ score. D) Abbreviated DotPlot with key activation markers and TCR signaling genes. The full DotPlot of phenotypic and activation markers is shown in supplemental figure S4.

While the majority of CD4+ T-cell subsets were relatively uncommitted to specific lineages, we found three more differentiated CD4⁺ populations: a TNFa-expressing activated CD4 T cell cluster (“activated CD4 T-cells”), a *ZBTB16* (PLZF)^+^ *KLRB1* (CD161)⁺ innate-like cluster (“ZBTB16+ CD4 T-cells”), and a FOXP3⁺ Treg cluster. Finally, TCR chain usage identified a substantial γδ T cell response. Among Vδ1 (*TRDV1*⁺) cells we found two distinct populations: an activated population expressing dendritic cell chemotactic ligands *XCL1*/2 and activation markers like *TNFRSF4* (CD134 / OX40) and *TNFRSF9* (CD137 / 4-1BB) (“activated Vδ1”) and one with a relatively more quiescent, *TCF7*-high state, expressing features associated with regulatory Vδ1 cells, including *NT5E* (CD73) and *IL10* (“quiescent Vδ1”). We also identified a Vγ9Vδ2 (*TRDV2*⁺ *TRGV9*⁺) cluster that displayed a pronounced cytotoxic phenotype (e.g. *GZMB*, *PRF1*) (“cytotoxic Vγ9Vδ2”). Lastly, we identified an additional cytotoxic *CXCR3+* γδ population with ambiguous expression of δV genes, precluding confident assignment to either Vδ1 or Vδ2 (“CXCR3+ cytotoxic γδ”).

The least differentiated cells retained CCR7 and were marked by immediate-early AP-1 genes (e.g. *BATF*), (“early-activated CD4⁺”). Along this activation gradient, cells progressively engaged distinct metabolic and stress programs: a glycolytic/hypoxic state (marked by *LDHA*, *ENO1*, *P4HA1*), an ER-stress/secretory state driven by the unfolded protein response (*XBP1*, *HSPA5*), a proteostasis state combining proteasome subunits with the integrated stress response (*PSMB6*, *PPP1R15A*) and, lastly, a pronounced biosynthetic signature indicative of proliferating cells (*DDX21*, *NME1*). Together, these clusters describe a trajectory of CD4⁺ T-cell activation: from early-activated cell states, through metabolic commitment and biosynthetic expansion, to full effector differentiation. Finally, we identified only one population of CD8+ T cells, with a quiescent, naïve-like phenotype devoid of effector or lineage-commitment genes.

To further delineate activated T-cells into those responding strongly to PfME antigen engagement, rather than simply bystander activation, we used a gene module approach, aggregating the expression of a combination of activation-induced markers and TCR-signaling components (see full list in methods) into an AIM score (**Fig 1C**). We identified four cell populations with the strongest evidence of Pf-responsiveness. Two populations were marked by an exceptionally high AIM score, the clusters we annotated as “activated CD4” and “activated Vδ1”. They expressed not only a breadth of costimulatory and type-I interferon-stimulated genes (*ICOS*, *IRF4*, *TNFRSF9* (CD137), *IL2RA* (CD25), *NFKB1*, *MIR155HG*), but also members of the Nur77 family, *NR4A1, NR4A2* and *NR4A3* supporting recent TCR engagement. In comparison, the “ZBTB16+ CD4” and “cytotoxic Vγ9Vδ2” populations had comparatively lower AIM scores and Nur77 expression but expressed higher levels of a large variety of activation-induced makers than the remaining clusters, including *CD69, CD40LG*, *ICOS*, *TNFRSF4* (CD134) and *TNFRSF18* (GITR). Further in support of functional TCR signaling among the ZBTB16+ CD4 population, we found evidence of emerging Th17 polarization: expression of *RORC* (RORγt) and *CCR6,* though we did not detect significant expression of IL17 genes in our data (**Fig. S5A**). The cytotoxic Vγ9Vδ2 cluster further distinguished itself by expression of *FCGR3A* (CD16a), as well as a suite of exhaustion markers, including *TIGIT, HAVCR2* (Tim3) and *ENTPD1* (CD39) (**Fig. S5B**), a phenotype suggestive of prior antigen exposure. We therefore considered these four populations (activated CD4, activated Vδ1, ZBTB16+ CD4 and cytotoxic Vγ9Vδ2 T cells) as demonstrating the highest degree of Pf-responsiveness and refer to their combined frequency as “total Pf-responsive T-cells”. Together these data suggest that PfME stimulation elicited broad activation of cord blood T-cells, dominated by CD4⁺ and γδ T cells with costimulatory and cytotoxic programs.

When assessed by maternal malaria status, the frequency of total Pf-responsive T cells, as well as each subset did not vary by placental malaria status (**Fig. S6**). In contrast, increasing number of maternal peripheral malaria infections was associated with significantly increased frequency of ZBTB16+ CD4 T cells (1.89% per infection, p=0.04, **Fig. S7**). Conversely, there was a trend towards reduced early activated CD4, the least differentiated CD4 T cell population, with increased maternal peripheral malaria infections, though this was non-significant (-6.8% per infection, p=0.056). Together, these results are consistent with more memory-like and fewer naïve-like CD4 T-cells in cord blood responding to PfME stimulation following maternal peripheral malaria exposure.

### Maternal microchimerism is associated with attenuated PIP

We next sought to determine the maternal or fetal origin of Pf-responsive T-cells. Using the approach outlined above, we identified 25 cells from six of the infants as MMc, however only 11 of these cells from five infants were retained following scRNAseq QC (**Fig 2A**). These 11 cells were broadly distributed across T-cell clusters, including early activated, proliferating, UPR/secretory, and glycolytic/hypoxic CD4 T cells; cytotoxic Vγ9δ2 T cells; and CD8 T cells. Thus, despite their diversity, the frequency of MMc amongst activated T-cells sorted for scRNAseq was low, suggesting that maternal origin cells were not a major contributor to the Pf-specific blood stage responses we identified in CBMC.

**Figure 2.**
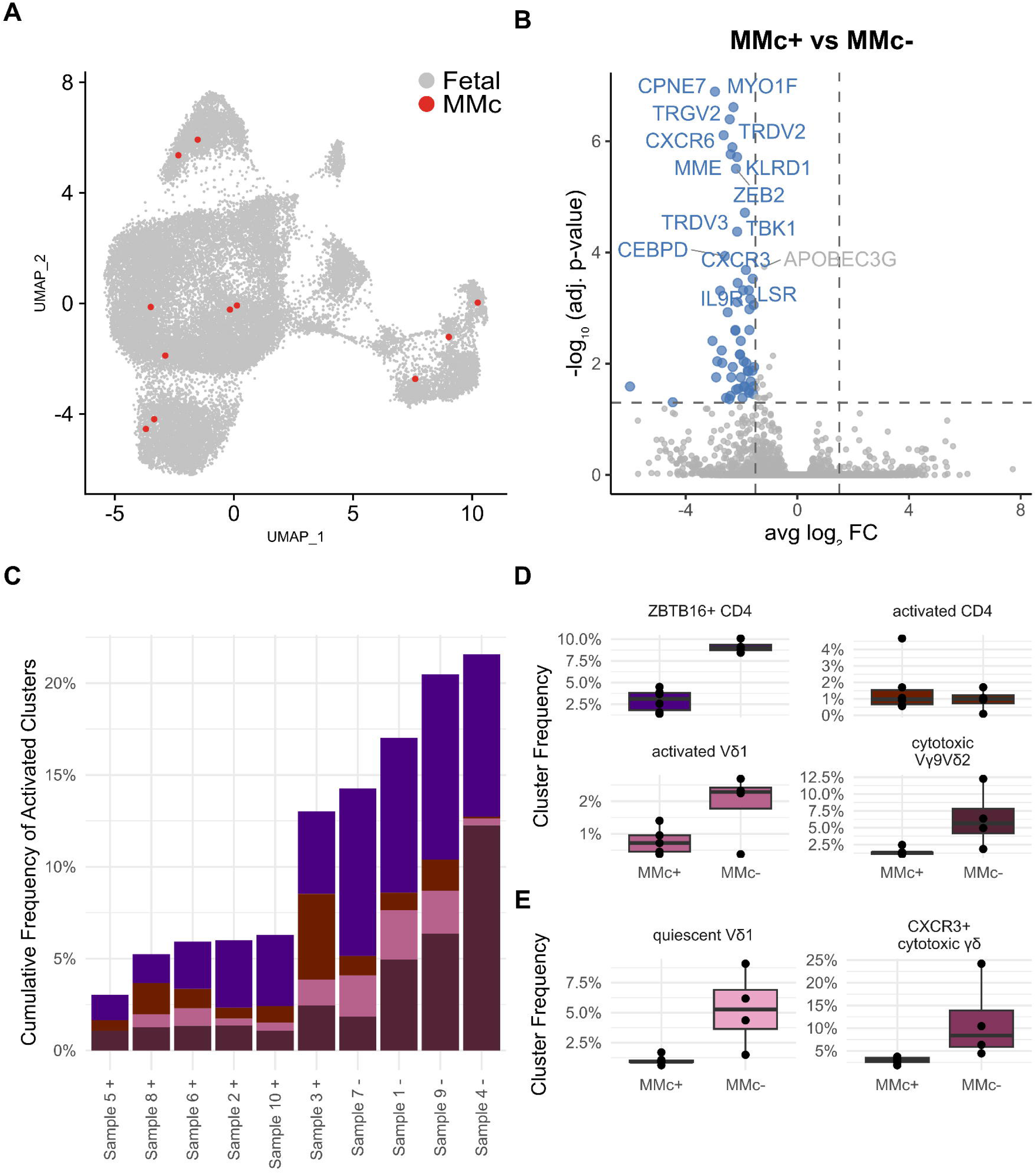
Maternal cells are detectable in Pf-responsive cord blood T cells and are associated with the frequency of activated cells. A) UMAP showing fetal cells in gray and high-confidence MMc+ cells in red. B) Volcano plot of pseudobulk differential gene expression results comparing all T cells from MMc+ samples with MMc-samples. C) Cumulative frequency and phenotypic composition of activated cells in each sample, colored identically to D) Frequency of all four activated T cell clusters. E) Additional clusters differentially abundant MMc+ and MMc-samples. Remaining data shown in figure S8.

To test whether the presence or absence of detectable MMc among activated T-cells impacted the transcriptional state of fetal cells, we performed pseudobulk differential gene expression. To account for the various ways in which maternal cells may impact the response to PfME, we performed differential gene expression between experimental MMc+ and MMc-samples in three ways: combining expression profiles of all T-cells, all activated T-cells with an AIM score >= 0.04, or all cells from the highly activated clusters (activated CD4, activated Vδ1, ZBTB16+ CD4, and cytotoxic Vγ9δ2). Results from all three analyses were similar, showing few differentially expressed genes (70 among all T-cells, 10 among all activated cells and 6 among all activated clusters). Among all T-cells, the 70 genes were comprised largely of innate-like T cell transcripts, including γ and δ vdj TCR genes, Killer Cell Lectin Like Receptors (*KLRD1, KLRC2)* as well as a variety of genes related to cytotoxicity, *PRF1*, *NKG7*, *EOMES*, *ZEB2* and *ZBTB16* (**Fig 2B**). The low number of differentially expressed genes and the co-regulation of gene sets related to innate-like T cell function is consistent with differences in cell frequency, rather than cell state, driving the differences in pseudobulk associated with MMc. This suggests that the key driver of differential T-cell responses associated with MMc status is differences in cell frequency, rather than transcriptional differences.

To next investigate our hypothesis that MMc may influence or “educate” the quality of the fetal response to Pf, we assessed whether the frequency of total Pf-responsive cells or individual subsets varied in MMc+ vs MMc-CBMC. We surmised that MMc could influence outcome even if the MMc cells did not survive scRNAseq QC, so we assigned experimental MMc status based on the distribution of the initial 25 MMc cells, resulting in four infants who were experimental MMc- and six infants who were experimental MMc+.

MMc+ samples had, on average, significantly fewer total Pf-responsive T-cells of sorted T-cells, relative to MMc-samples (6.6% vs. 18.3%, p<0.001) (**Fig 2C**). Specifically, the frequency of Pf-responsive ZBTB16+ CD4 T cells (2.9% vs. 9.11%, p<0.001), Vδ1+ T cells (0.7% vs. 1.9%, p=0.03), and cytotoxic Vγ9δ2+ T cells (1.4% vs. 6.4%, p=0.02) were all lower in MMc+ vs MMc-samples on average (**Fig 2D**). By comparison, the frequency of Pf-responsive activated CD4 T cells did not vary (1.6% vs. 1.0%, p=0.5). These differences persisted after adjustment for maternal placental and peripheral malaria, suggesting that the association was independent of maternal malaria status. Two additional subsets with lower Pf-responsiveness (quiescent Vδ1 and CXCR3+ cytotoxic γδ) were also significantly lower in MMc+ vs MMc-samples (**Fig 2E**).

### MMc and Pf-responsive T-cell populations display differential association with risk of malaria outcomes during infancy

To understand the clinical implications of these observations, we next assessed whether either the presence of MMc or the quality of PIP influenced clinical malaria outcomes during early childhood. Among 8 infants followed for at least 4 weeks, there were 102 clinical visits, 15 (15%) of which were incident parasitemias. Among these incident parasitemias, 6 (40%) were associated with fever. Experimental MMc+ status was associated with protection from incident parasitemia (AOR=0.47, p=0.03) and febrile malaria when infected (AOR=0.10, p=0.04) (**Fig 3**). In contrast to MMc, the total frequency of Pf-responsive T cells was associated with increased incident parasitemia (AOR=1.08 per 1% increase, p<0.001). When the frequency of the four Pf-responsive subsets were modeled together, ZBTB16+ CD4 T-cells (AOR=2.01, p<0.001) and activated CD4 T cells (AOR=2.23, p<0.001) were associated with increased risk of incident parasitemia. In contrast, Vδ1 T cells were associated with a trend toward protection (AOR=0.38, p=0.09) and Vγ9δ2 T cells had no strong association (AOR=0.87, p=0.2). Finally, when MMc was added to the multivariate model, both the protective association of MMc (AOR=0.27, p=0.68) and the risk association of ZBTB16+ were attenuated (AOR=1.39, p=0.59), suggesting that they operate through a shared pathway; whereas, the associations with activated T cells (AOR=1.97, p=0.005) and Vδ1 T cells (AOR=0.45, p=0.1) were sustained.

**Figure 3.**
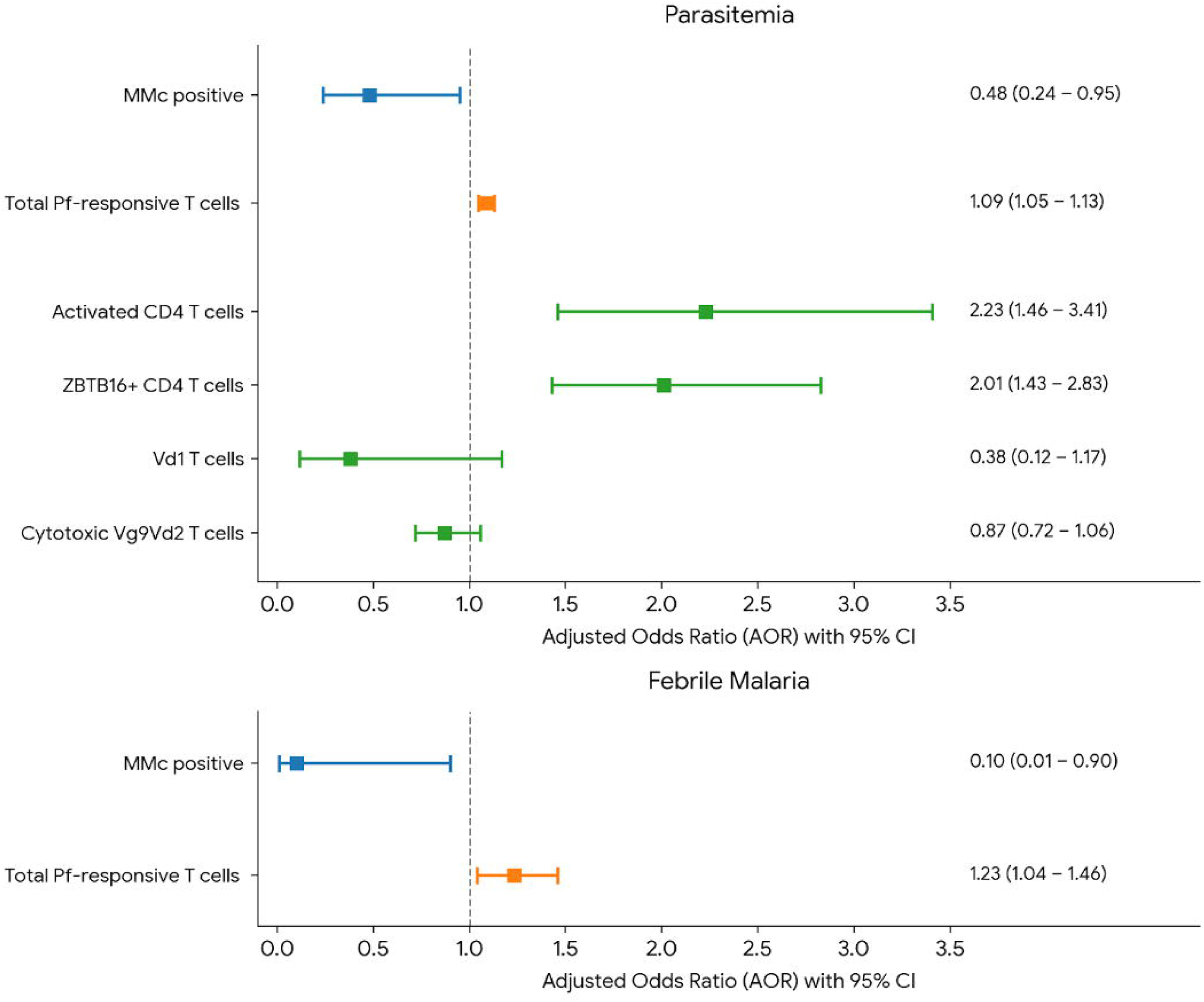
Association between maternal microchimerism, Pf-specific prenatal immune priming, and malaria outcomes. Malaria outcomes modeled in children followed for at least 4 weeks (n=8) using binomial GEE models with an independent correlation structure and robust standard errors. Impact of MMc (blue) modeled in univariate analysis, impact of total Pf-responsive T cells (orange) modeled in bivariate analysis adjusted for frequency of CD69+ cells of sorted CD3 T-cells, and impact of Pf-responsive T cell subsets (green) modeled in multivariate analysis also adjusted for frequency of CD69+ cells of sorted CD3 T-cells. A) Adjusted odds of parasitemia. B) Adjusted odds of febrile malaria in the setting of parasitemia.

Similarly, the frequency of total Pf-specific T cells was associated with increased risk of febrile malaria (AOR=1.23, p=0.01). Unfortunately, there were not sufficient outcomes to achieve model convergence when all subsets were modeled together, or when MMc was added, however, in univariate analysis, ZBTB16+ T-cells were associated with increased risk (AOR=1.69, p<0.001), whereas activated CD4 T cells were associated with decreased risk (AOR=0.42, p<0.001). Because ZBTB16+ T cells are easily bystander activated, we additionally assessed the association between ZBTB16+ T-cell frequency and non-malarial fever. Consistent with our malaria observations, MMc was protective from non-malaria fever (AOR=0.17, p=0.05), whereas ZBTB16+ T cells were associated with increased risk of non-malarial fever (AOR=1.34, p=0.005). No other Pf-responsive T-cell subset was associated with risk of non-malarial fever. Taken together, these data are consistent with a model in which the impact of MMc on clinical outcomes may be mediated through regulation of ZBTB16+ cells responses in cord blood, where these populations are associated with increased risk of malaria and non-malarial fever during infancy (**Fig 4**).

## DISCUSSION

Using single cell approaches, we identified four distinct subsets of T-cells in Malian cord blood responsive to blood stage Pf antigens: two CD4 and two γδ T-cell populations. Our “activated CD4 T-cells” demonstrated recent TCR engagement, evidenced by several activation markers and by NR4A1 (encoding Nur77), a specific marker of antigen receptor signaling. These cells lacked lineage commitment markers but expressed TNF, potentially indicative of a central memory fate, although the short stimulation may have precluded robust induction of lineage markers in previously naïve cells. This cell subset was associated with increased risk of parasitemia during infancy, which may reflect the dissociation between antigen-specificity and functional protection, and in particular the inability to robustly produce IFN. This is also in contrast to the association recently reported between frequency of Pf-responsive CD4 T-cells and protection from malaria outcomes (17). However, that study measured proliferation of total CD4 T-cells, so our transcription-based approach may have resolved a distinct subset.

Our discovery of *Pf-*responsive *ZBTB16*+ (PLZF+) CD4 T cells that co-express *KLRB1* (CD161) is notable for several reasons. CD161+ CD4 T-cells have been shown to be increased in adults following vaccination with irradiated *P. falciparum* sporozoites and were associated with protection (33). CD161+ CD4 T cells were also associated with improved parasite control in human challenge studies with semi-immune adults (34). In contrast, we found these cells to be associated with greater susceptibility to blood stage parasitemia during infancy. This discrepancy may be explained by age-associated differences in CD161+ T cells: *ZBTB16* expression is largely absent in peripheral T-cells in both older children and adults, while it is widespread in many fetal tissues (35), conferring distinct functionality to fetal CD161+ T cells. In our study, ZBTB16+ cells showed lower direct TCR engagement, suggesting activation by cytokines rather than antigen, consistent with reports that cord blood ZBTB16+ CD4 T cells are easily activated and rapidly amplify inflammatory responses, recruiting neutrophils and macrophages (36). Alternatively, this may reflect KLRB1 itself, a negative regulator of TCR signaling (35). Nonetheless, these cells showed emerging Th17 polarization (*RORC*, *CCR6*) (37). Th17 responses to *P. falciparum* are unlikely to be functionally beneficial and may thus explain their association with increased susceptibility to parasitemia in our data.

We also found two Vδ1 T cell populations, one highly activated and one more quiescent. The activated population showed a trend toward protection from parasitemia, consistent with reports associating Vδ1 expansion with asymptomatic malaria in Ghanaian children (38) and with natural infection and controlled human malaria infection (39). The quiescent population may perform regulatory functions: these cells expressed *IL10* and *NT5E* (CD73), which generates immunosuppressive adenosine (40). Lastly, we identified cytotoxic Vδ2 T cells, which have been well described as malaria responsive (41), and appeared to be TCR engaged. These cells are notable as they were distinct from other cells in our data through expression of a suite of exhaustion markers, including *HAVCR2* [Tim3], *TIGIT* and *ENTPD1* [CD39]. This phenotype is associated with sustained TCR signaling and thus unlikely to be induced following an 18h stimulation of previously naïve cells. Likewise, their expression of *FCGR3A* (CD16) implies previous encounter with antigen. Together, their phenotype is consistent with extensive previous exposure to antigen, resulting in a terminally differentiated cytotoxic effector γδ T cell population.

Our epidemiologic analysis of the screened samples supported our prior observation that MMc is associated with increased risk of parasitemia but decreased risk of febrile malaria, thus we next queried the origin of the Pf-responsive cells. We identified few MMc T cells in stimulated CBMC, but their frequency (∼10^-3^) was substantially higher than MMc typically detected in cord blood (∼10^-5^-10^-4^), suggesting relative concentration of Pf-specific MMc, and those present were phenotypically diverse. Most notably, MMc in stimulated CBMC was strongly associated with PIP, in particular a lower frequency of TNF-producing ZBTB16+ CD4 T cells. This is consistent with mechanistic mouse studies in which antigen-specific MMc regulate inflammatory responses to matched pathogen challenge, reducing *ZBTB16* expression, inflammatory infiltration and immunopathology (Toure, unpublished).

Supporting the clinical relevance of these observations, we identified an association between MMc in our sorted CBMC and risk of both parasitemia and febrile malaria during early childhood. The observation of protection from parasitemia contrasts with our screened MMc results, from both Tanzania (22) and this cohort, but this may reflect differences in the MMc identified through the two approaches. Specifically, in our qPCR assays we detect all MMc in whole cord blood, reflecting a variety of immune cell types as well as both antigen-specific and non-specific T cell populations. In contrast, MMc captured in our scRNAseq data are restricted to T cells that had high expression of CD69 following PfME stimulation.

Taken together, our data support a model in which the mother develops Pf-specific T cells before and during pregnancy that may be transferred to the fetus *in utero*, while the fetus independently generates a diverse, multi-lineage response to Pf antigens. MMc appears to “educate” this response, restraining highly inflammatory *ZBTB16*+ CD4 T cells, which may ultimately reduce the risk of febrile malaria in infancy. Thus, although the fetus acquires few MMc, they may have a disproportionate impact by setting the tone of the offspring immune response.

Our study has several limitations. Although we screened a large cohort, we described PIP at the single cell level in only ten infants due to resource limitations; nonetheless, we identified clear evidence of PIP in all infants and strong associations with clinical outcomes. Second, PfME, although a complex blood stage antigen source, would not contain high levels of PfEMP1, and future work should target PfEMP1-directed responses. Finally, we cannot exclude a role for MMc in modulating responses at the time of acute Pf exposure, although we previously found few MMc during acute malaria (21), and in a South African cohort birth rather than concurrent MMc was the strongest determinant of BCG recall responses (27), suggesting MMc acts primarily on very early immune development.

In conclusion, we demonstrate for the first time the complexity of the early immune response to malaria, including both fetal priming and maternal regulation of this priming. These observations highlight the importance of intergenerational immune interactions in shaping offspring immunity. The consistent observation that MMc is associated with protection from infection, and in particular immunopathology, observed in our and others work, highlights the critical role that these cells play in offspring immunity.

