## Supplemental Materials for "Prenatal immune priming to *Plasmodium falciparum* is characterized by multi-lineage T-cell responses that are influenced by maternal microchimerism and predict malaria outcomes"

**Affiliations:**
^1^ Center for Global Infectious Disease Research, Seattle Children’s Research Institute, Seattle, WA, USA

^2^ Division of Infectious Diseases and Geographic Medicine, Stanford University School of Medicine, Stanford, CA, USA

^3^ International Center for Excellence in Research, Bamako, Mali

^4^ Laboratory of Malaria Immunology & Vaccinology, National Institute of Allergy and Infectious Disease

^5^ Department of Biomedical Sciences, Iowa State University, Iowa, USA

^6^ Department of Pediatrics, University of Washington School of Medicine, Seattle, USA

^7^ Department of Pediatrics, University of Maryland School of Medicine, Baltimore, USA

^8^ Department of Immunology, University of Washington, Seattle, USA

**Corresponding Author:**

Whitney E Harrington, MD PhD

1916 Boren Ave, Seattle, WA 98101

### Contents:

**Figure S1. Overview of the experimental approach to identify prenatal immune priming.**

**Figure S2** **Validation of approach to identify maternal microchimeric cells.**

**Figure S3. Association between screened MMc in cord blood and risk of malaria outcomes across early childhood.**

**Figure S4. DotPlot with all phenotypic, activation and functional genes considered for cluster identity assignment.**

**Figure S5. UMAPs with key exhaustion and Th17 markers.**

**Figure S6. T-cell cluster frequency stratified by placental malaria status.**

**Figure S7. T-cell cluster frequency stratified by malaria incidence during pregnancy.**

**Figure S8. T-cell cluster frequency stratified by experimental MMc status**

**A**

**B**

**C**

PfME

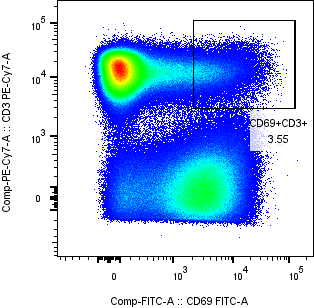

puRBCE

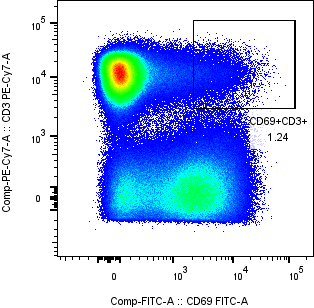

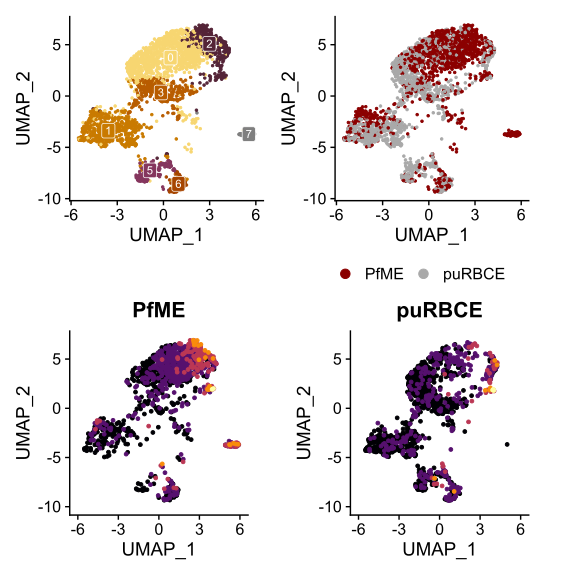

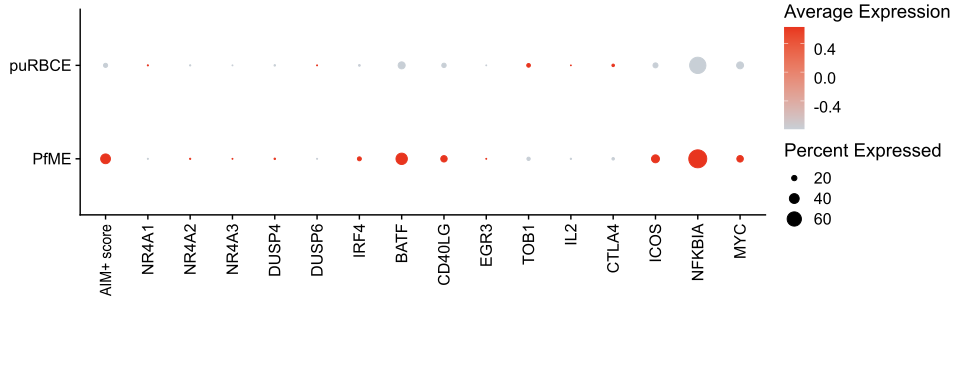

**D**

**Figure S1. Overview of the experimental approach to identify prenatal immune priming.** Validation of the approach using Pf-exposed adult PBMC stimulated with purified uninfected RBC extract (puRBCE) and *Plasmodium falciparum* merozoite extract (PfME). A) Sort gates of CD69+ CD3+ live single T cells used for subsequent scRNAseq with 10X. B) Assignment of cell identity using Louvain clustering. C) Discrimination of TCR-engaged cells from bystander activated cells using our AIM+ gene signature and key genes associated with T cell activation.

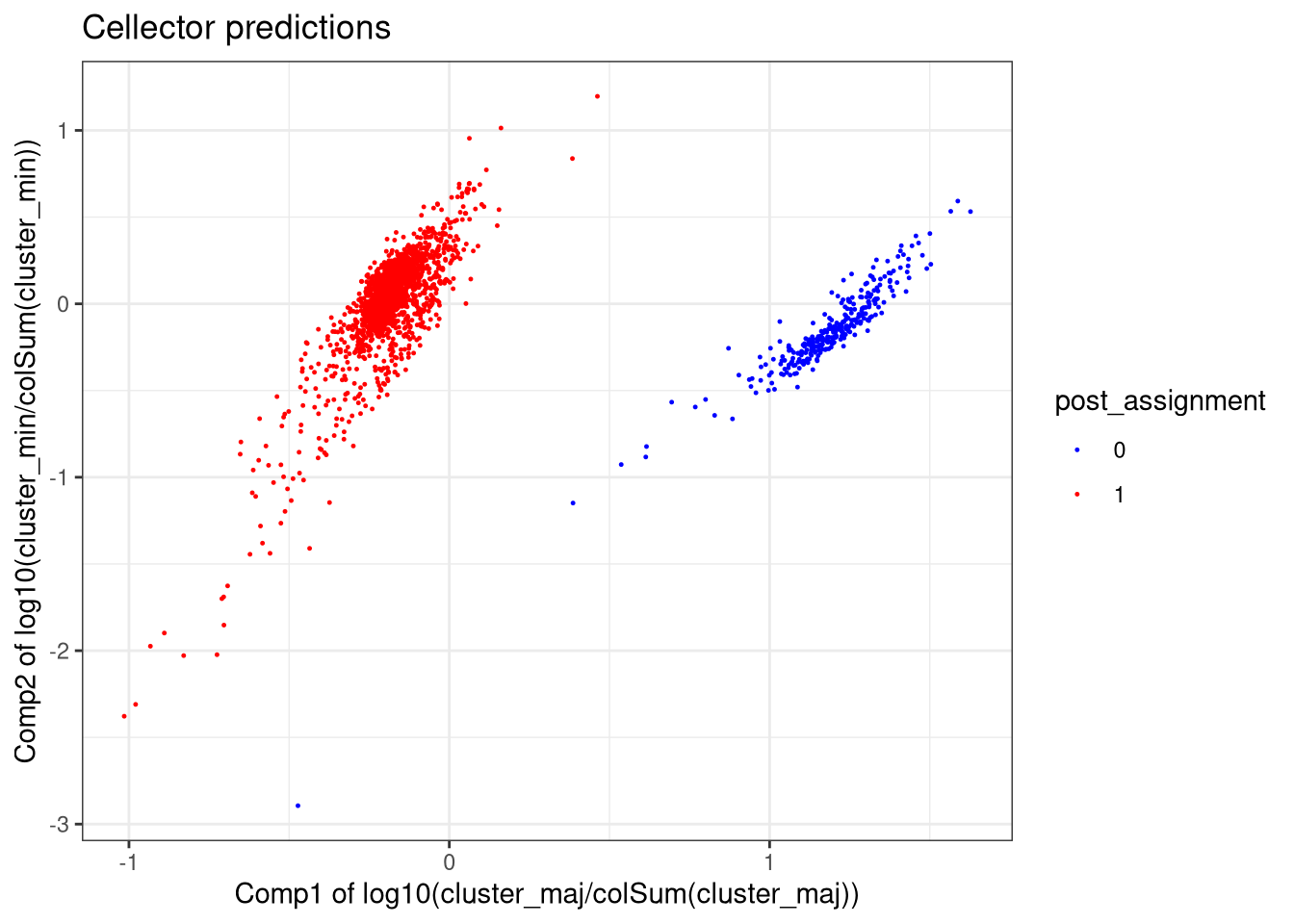

Cellector Assignment

**A**

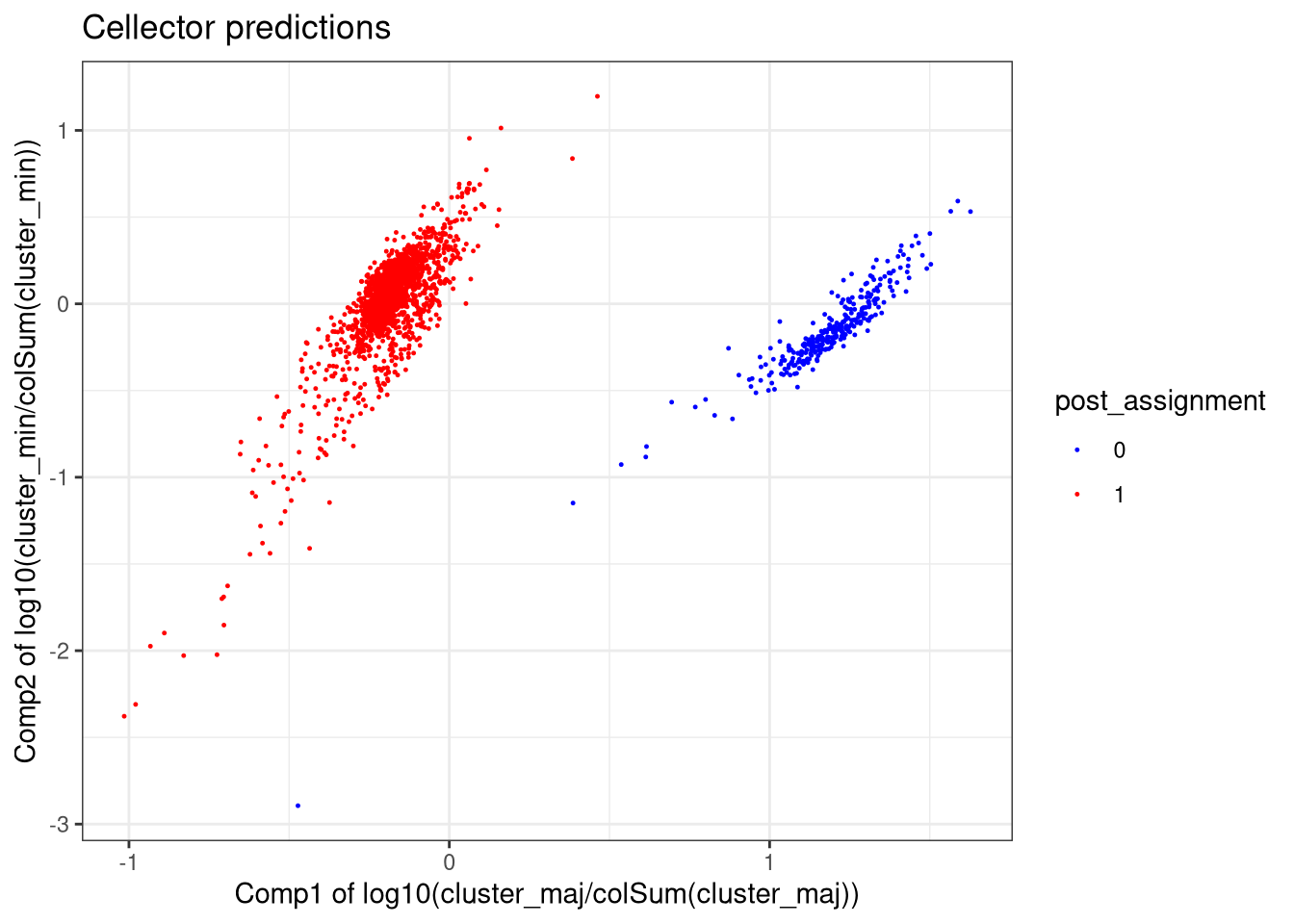

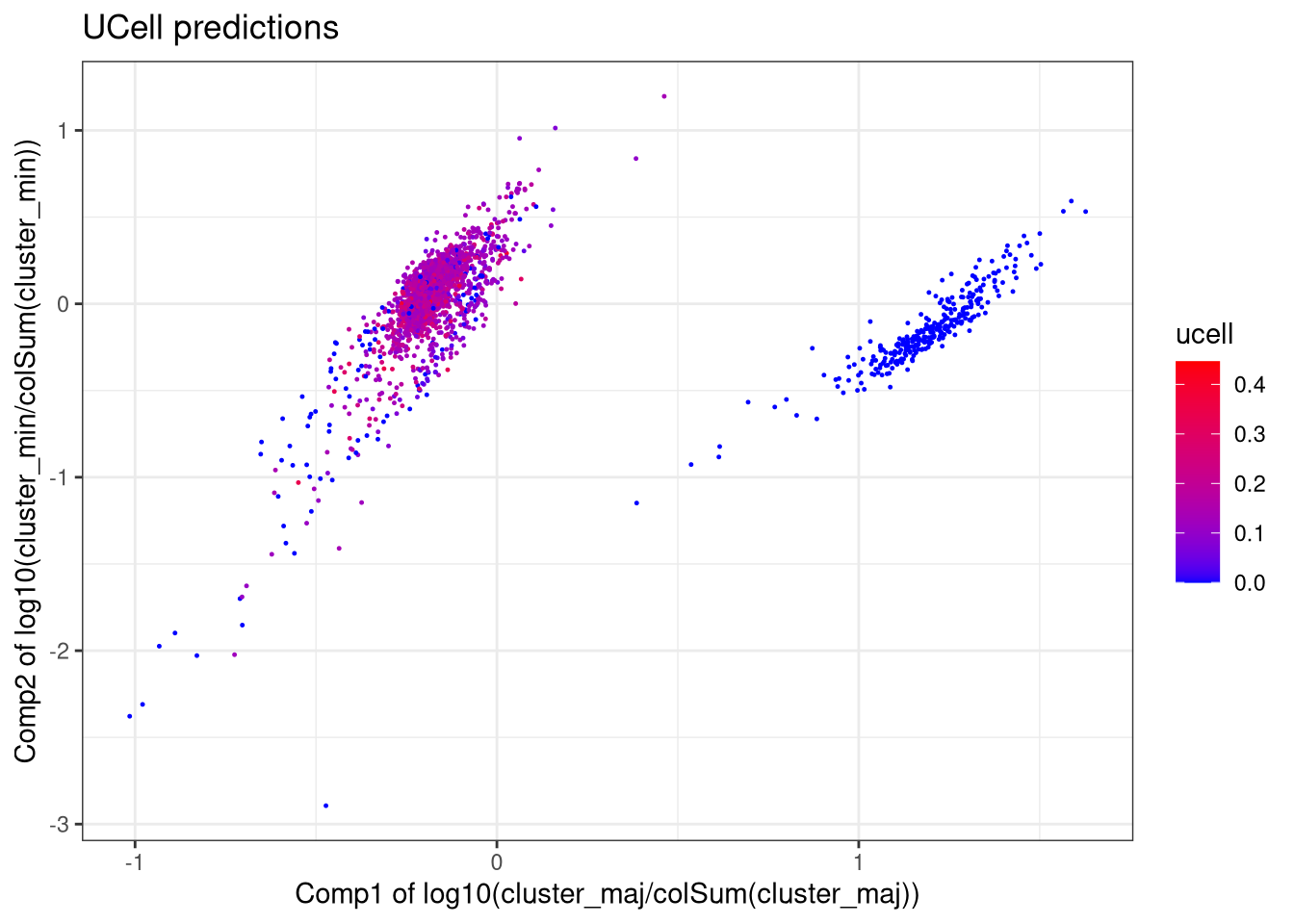

Y Chromosome Expression

**B**

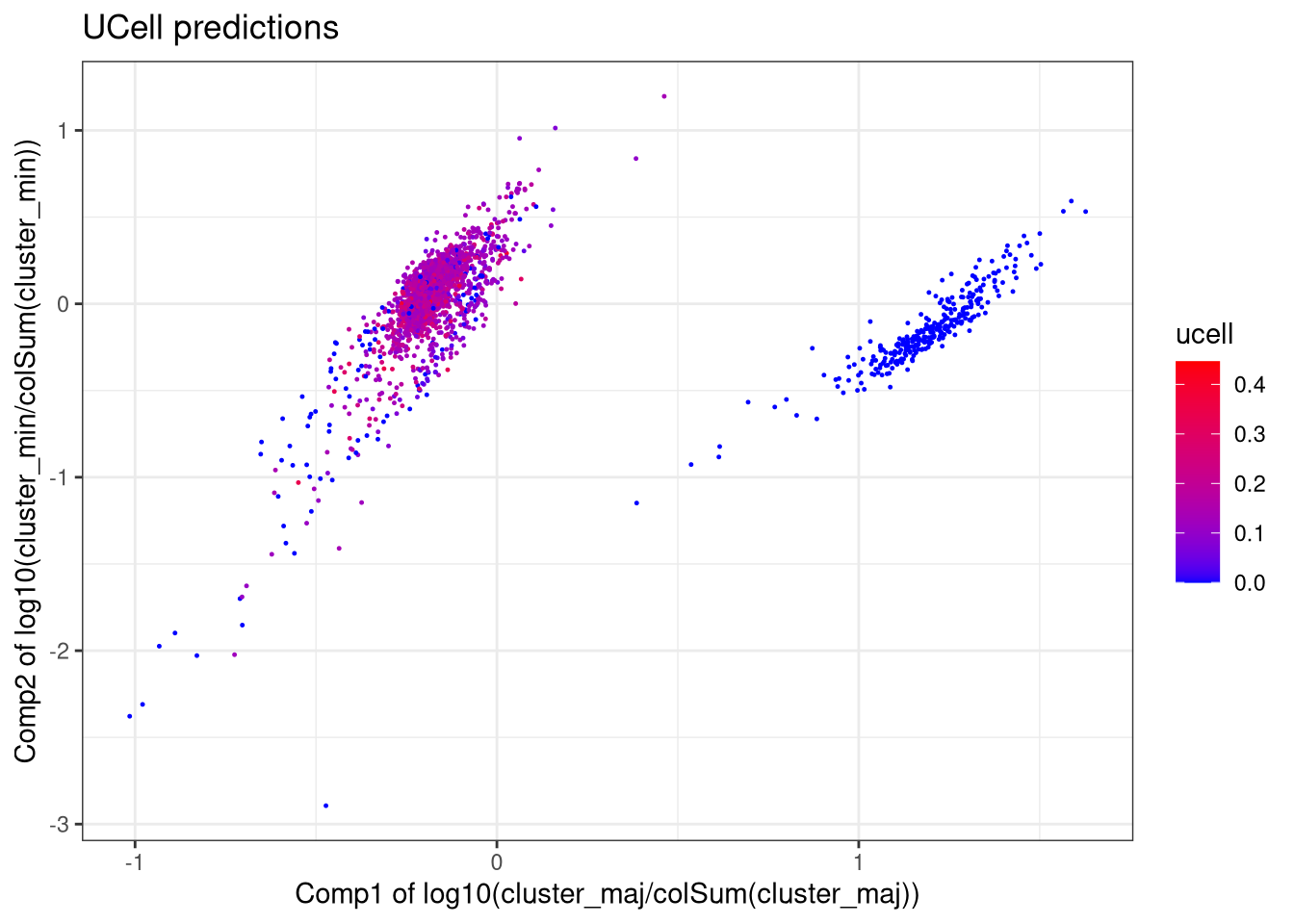

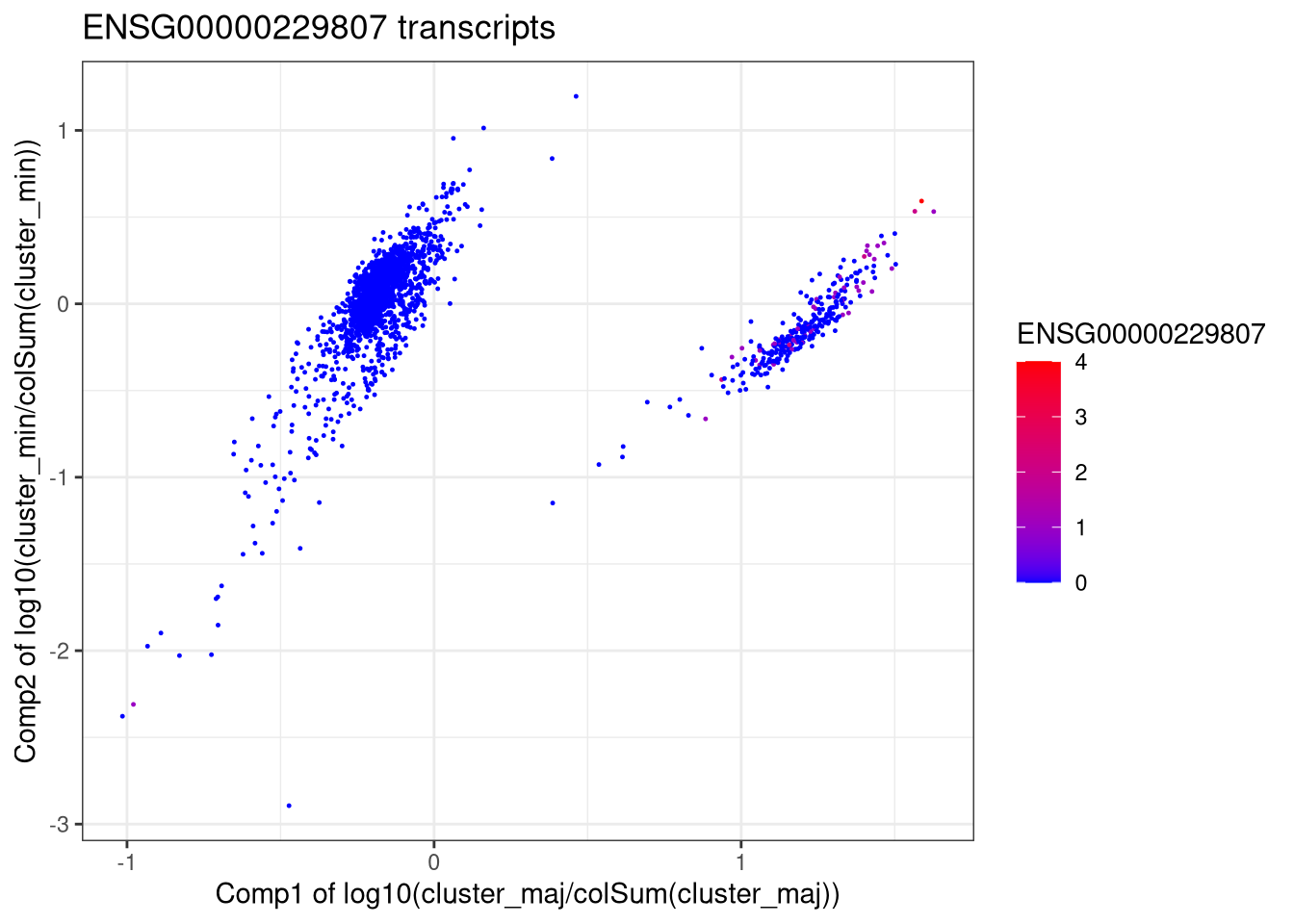

XIST Expression

**C**

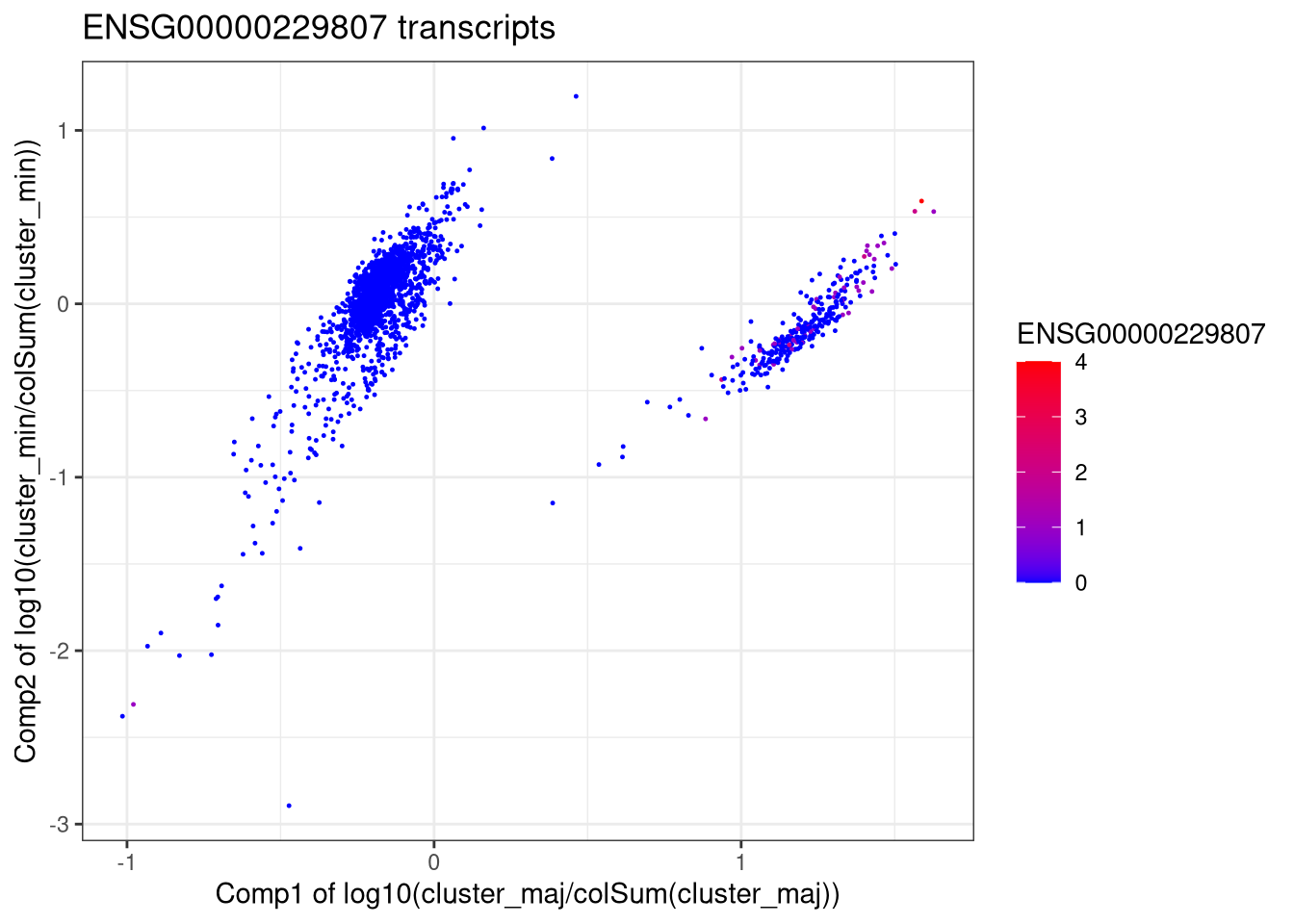

XIST

**Figure S2** **Validation of approach to identify maternal microchimeric cells.** PBMC from a female Pf-exposed donor were stimulated with PfME and CD69+ CD3+ T cells were added into unstimulated PBMC from a male donor in a ratio of 15% to 85%. 2,255 cells were sequenced with scRNAseq using the 10X platform. A) Cellector assigned identity as minority (0) or majority (1) population, plotted as negative log majority likelihood versus negative log minority likelihood. B) Overlay of integrated expression of seven genes from the Y chromosome (RPS4Y1, DDX3Y, KDM5D, EIF1AY, ZFY, UTY, and USP9Y) using Ucell. All cells with Y chromosome gene expression fell within the majority population. C) Overlay of XIST expression. All cells with XIST expression fell within the minority population.

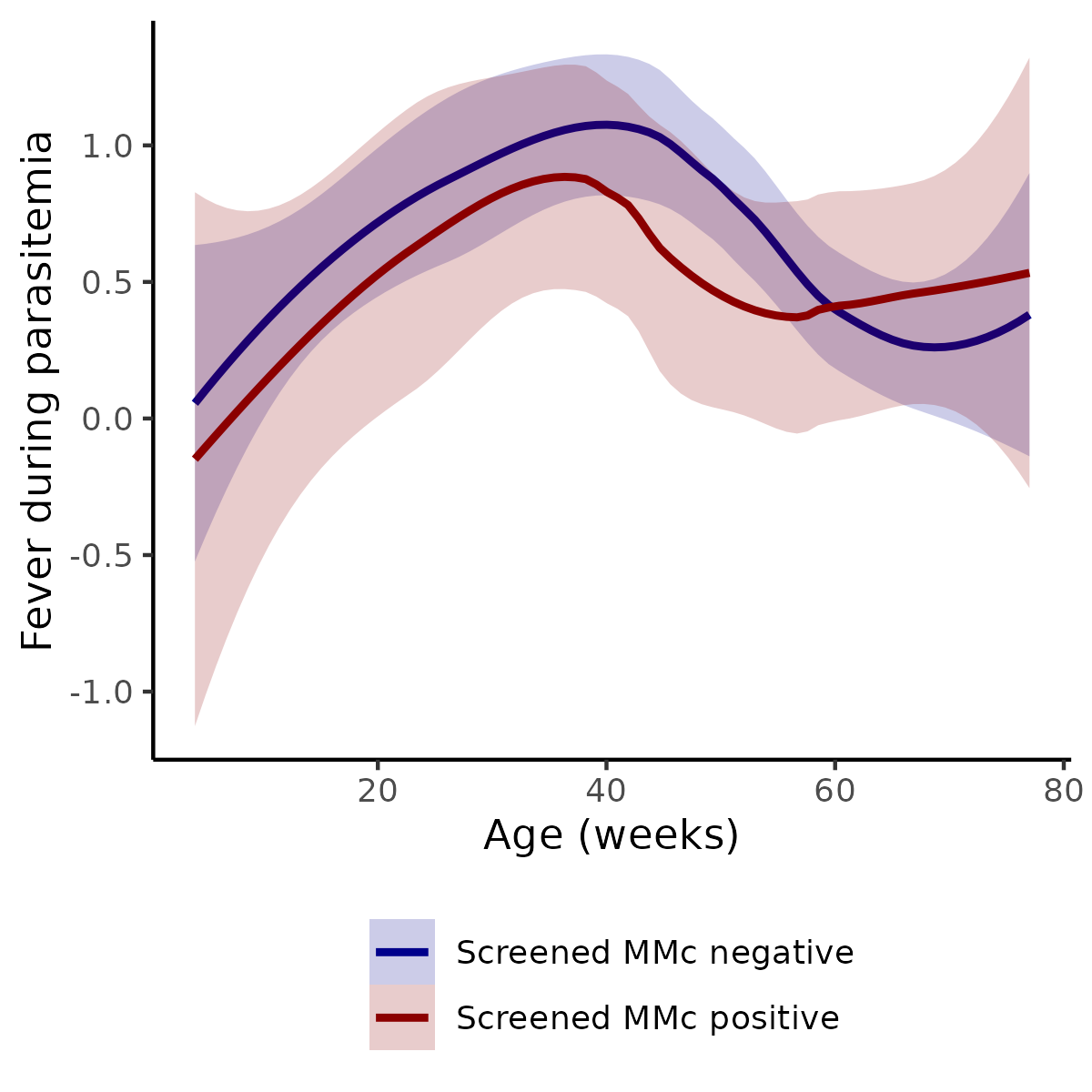

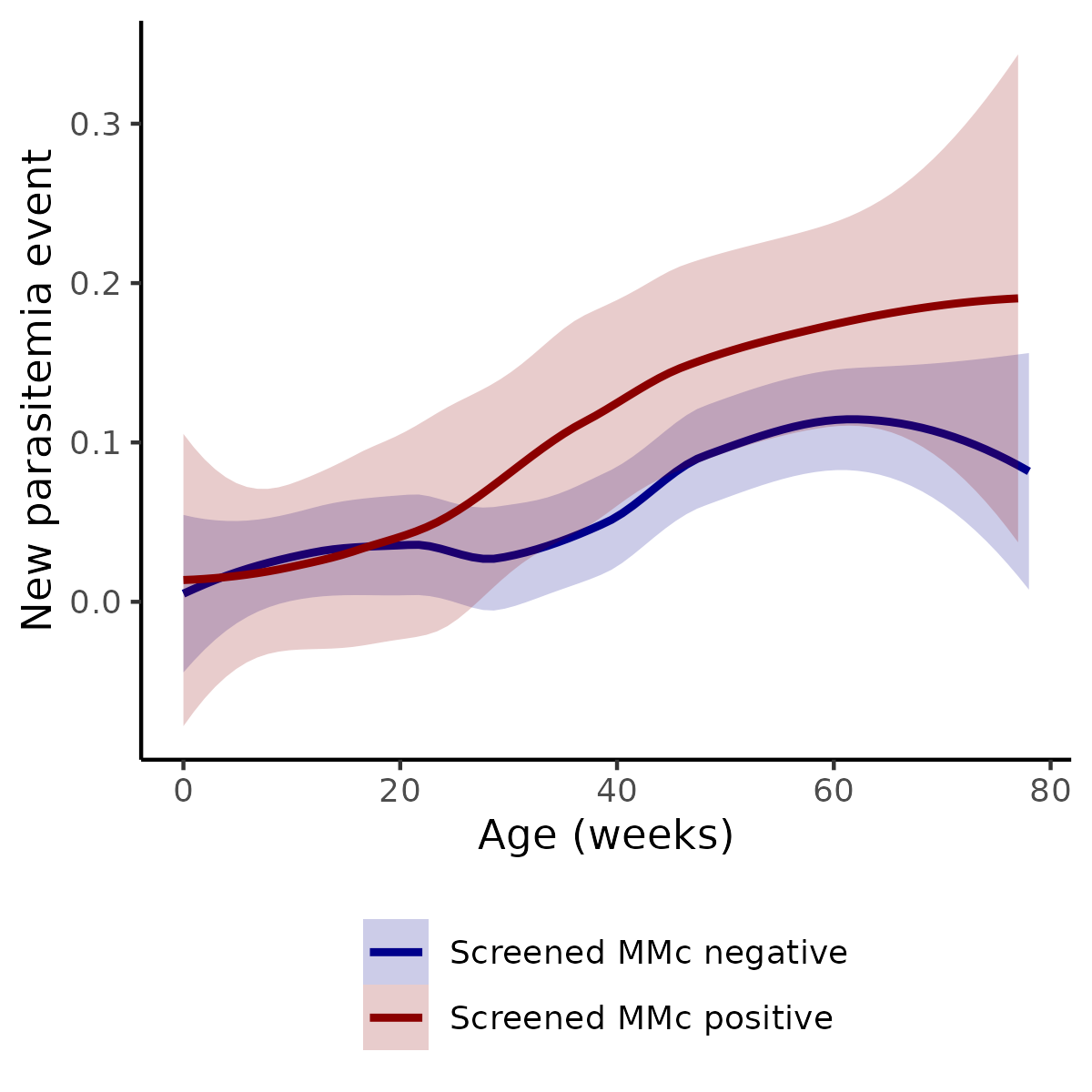

**A**

**B**

**Figure S3. Association between screened MMc in cord blood and risk of malaria outcomes across early childhood.** Outcomes modeled with data from 890 visits from 63 children followed for at least four weeks. A) Lowess curve of proportion of visits with incident parasitemia out of all visits stratified by cord blood MMc detection. B) Lowess curve of proportion of febrile malaria out of all parasitemia positive visits stratified by cord blood MMc detection.

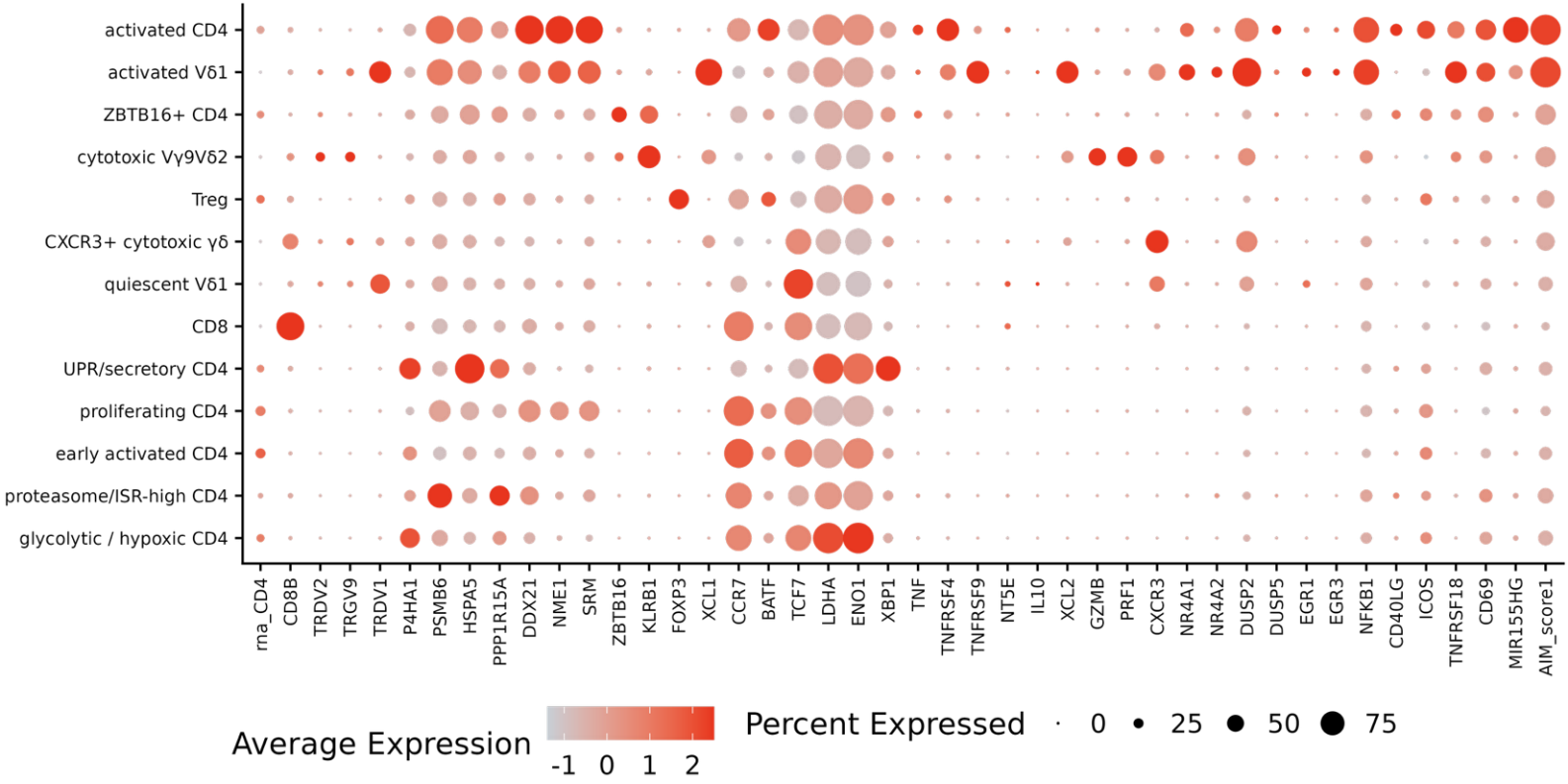

**Figure S4. DotPlot with all phenotypic, activation and functional genes considered for cluster identity assignment.**

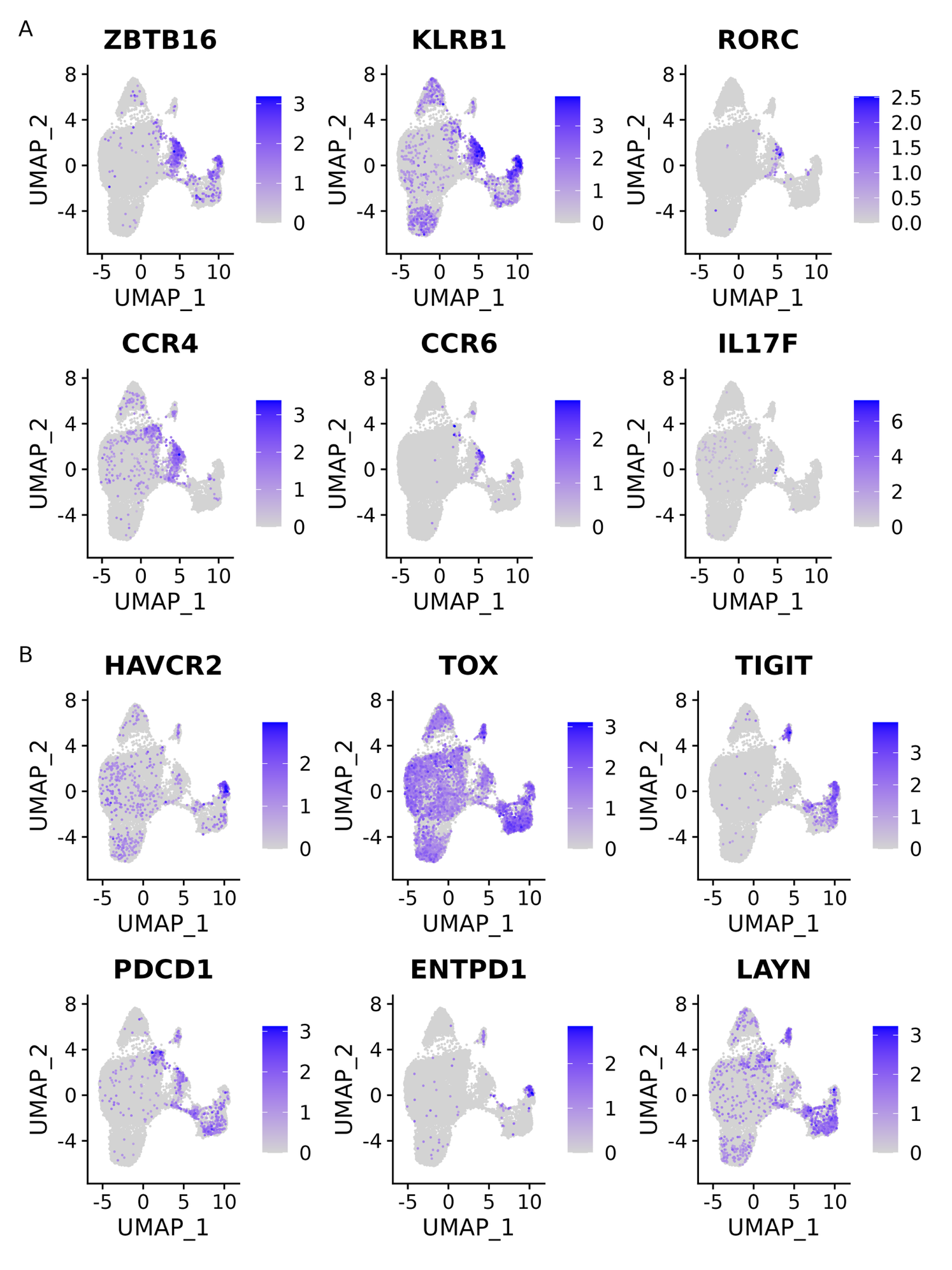

**Figure S5. UMAPs showing A) key Th17 and B) key exhaustion markers.**

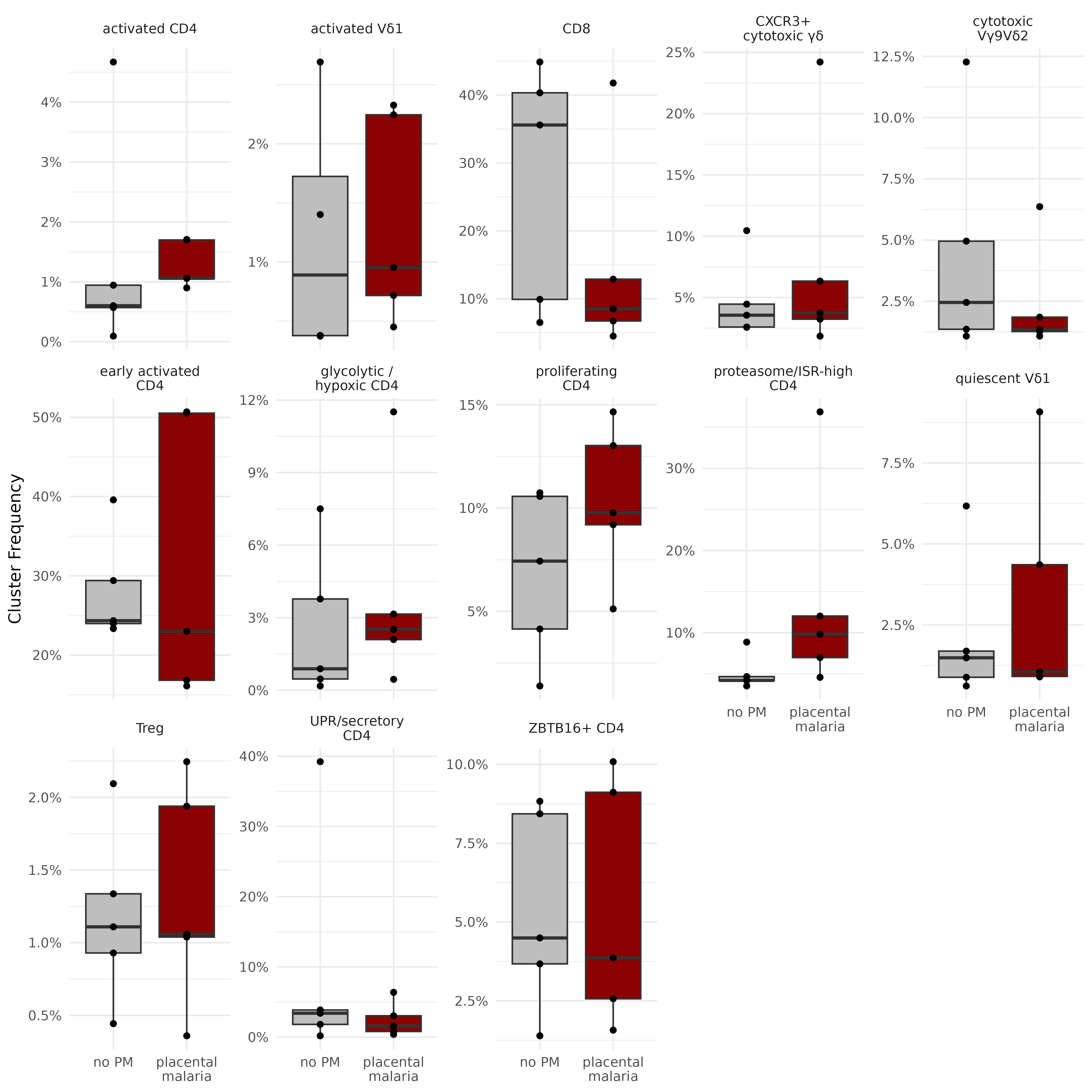

**Figure S6. T-cell cluster frequency stratified by placental malaria status.**

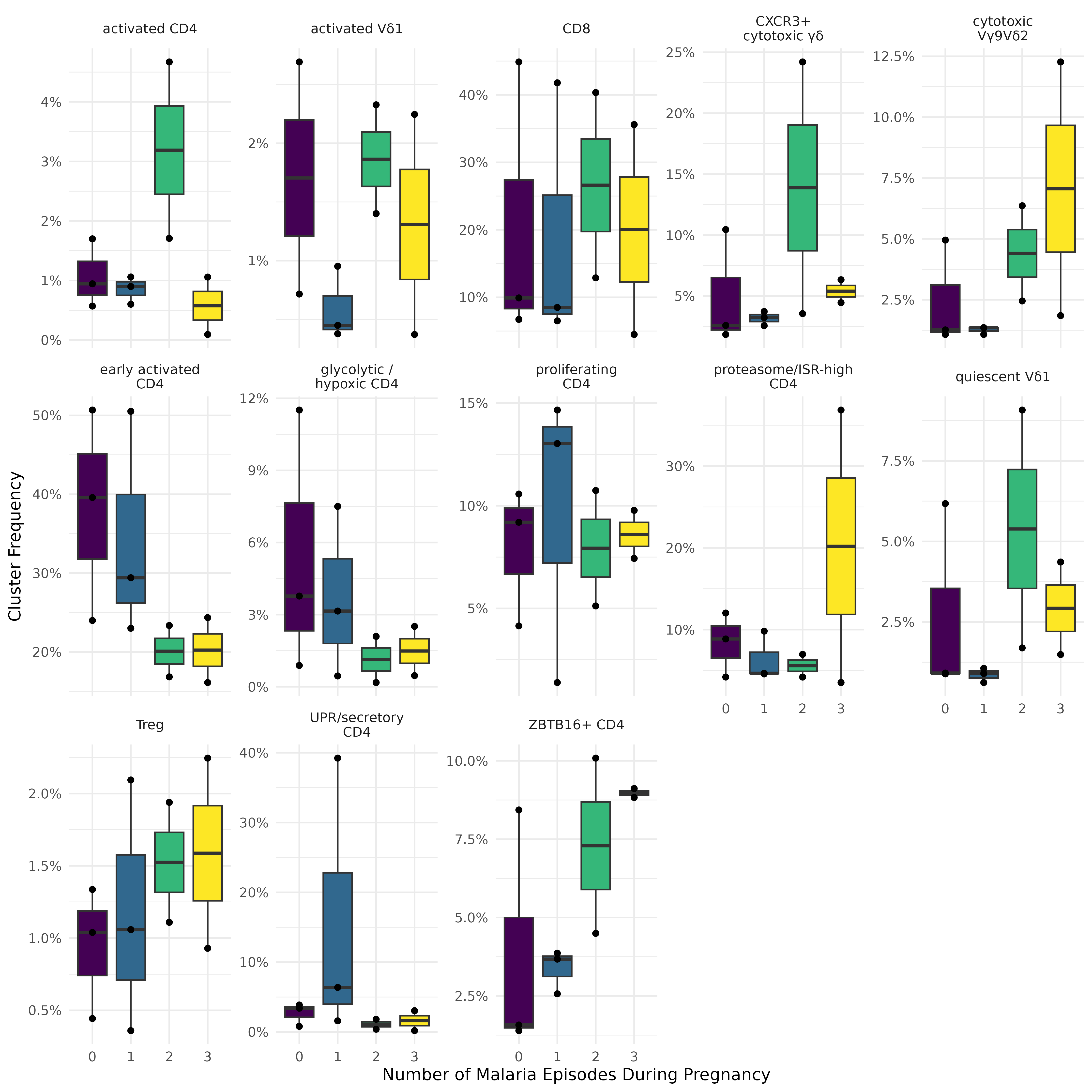

**Figure S7. T-cell cluster frequency stratified by malaria incidence during pregnancy.**

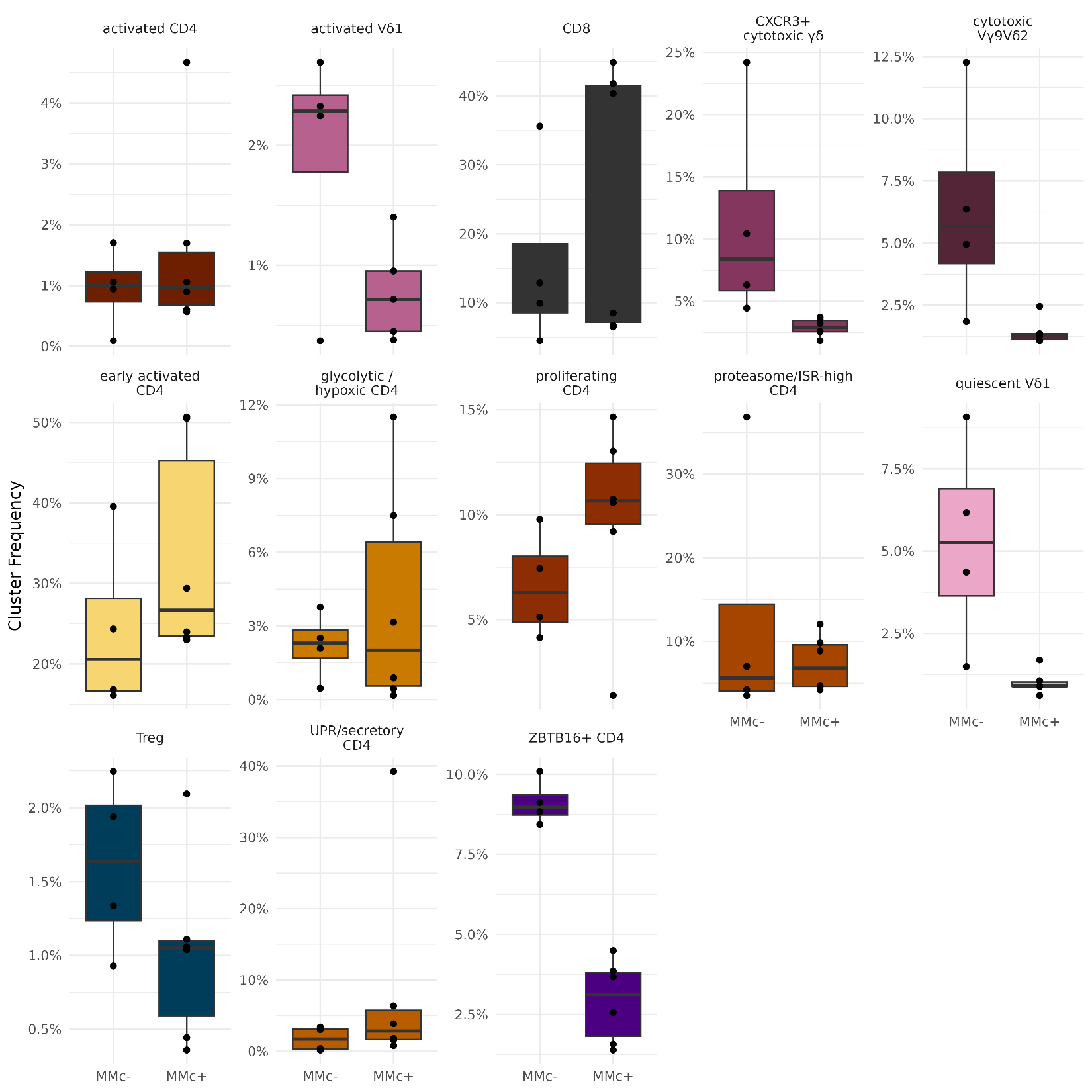

**Figure S8. T-cell cluster frequency stratified by empirical MMc status**
